# OneGenome-Rice (OGR): A genomic foundation model for rice

**DOI:** 10.64898/2026.04.21.719822

**Authors:** Bilian Qian, Chengwei Liang, Chao Qin, Chao Liu, Chunling Zhang, Chunyan Xu, Dongxu Li, Guirong Xue, Hang He, He Zhang, Huiying He, Duoyuan Chen, Jiwei Xu, Junyang Zhang, Jian Sun, Lianguang Shang, Jinling Jiang, Keke Xia, Liyuan Zhong, Ling-Ling Chen, Longjiang Fan, Longqi Liu, Mumu Qin, Qian Li, Rui Huang, Shenjun Zhu, Shengwei Ma, Shiping Liu, Shiyu Zhang, Shuai Fu, Tong Wei, Xiaolong Xu, Xinye Jia, Xun Xu, Yi Jing, Yu Xu, Yutao Bian, Yinuo Zhao, Yunlong Xue, Yafei Guo, Zhan Xiao, Zhaorong Li, Zhefu Li, Zhen Yue, Ziqing Deng

## Abstract

The transition of genomics to a predictive intelligence discipline is driven by the advent of genomic foundation models. While substantial progress has been observed in all-life and human-centric models, plant species, particularly for the staple crops, remains hindered by a lack of models. Here we introduce OneGenome-Rice (OGR), a rice (*Oryza sativa*) genomic foundation model pre-trained on a genomic dataset comprising 422 high-quality genomes of cultivated and wild rice. OGR is engineered upon a Mixture of Experts (MoE) transformer architecture with 1.25-billion parameters and supports an ultra-long context window of up to 1 million base (Mb) pairs at single-nucleotide resolution. A comprehensive benchmark demonstrated that OGR significantly outperforms existing state-of-the-art multi-plant or all-life genome models in 11 categories (e.g. motif identification, sweep detection, etc). We further demonstrated the utility of OGR in several downstream applications, such as *indica*-*japonica* subspecies introgression analysis, identification of agronomy trait-associated functional loci and prediction of gene expression from DNA sequences. These results establish OGR as a promising foundational computational infrastructure for rice functional genomics and precision breeding. The OGR and its fine-tuned models, including pretrained weights, training code and the rice genomic benchmark suit, have been fully opened.

## INTRODUCTION

The decoding of genomic sequences to understand the complex regulatory code of life is a cornerstone of modern quantitative biology. Genomic sequences encode a vast array of information, from local protein-coding instructions to distal regulatory grammars that orchestrate tissue-specific expression programs and developmental trajectories. In the agricultural sector, the ability to accurately predict the functional consequences of genetic variation is essential for addressing the dual challenges of global food security and climate change. Rice (*Oryza sativa*) genomic landscape is defined by its two major subspecies, *indica* and *japonica*, which diverged significantly and were subjected to independent domestication processes (Wing et al., 2018). Rice sustains more than half of the world’s population, serves as the premier model for cereal genomics due to its relatively compact genome and its pivotal role in the evolutionary history of the grass family (Huang et al., 2012; Wu et al., 2023; Fornasiero et al., 2025; Guo et al., 2025).

In recent years, deep learning-based genomic foundation models have emerged as a transformative technology (Boshar et al., 2025; Brixi et al., 2026; Avsec et al., 2026). These models leverage large-scale self-supervised pre-training on unlabeled DNA to learn the grammar and semantics of the genome, which can then be adapted to various downstream tasks with minimal task-specific data. While human-centric models like AlphaGenome, Genos, and multi-species models like Evo2, NTv3, and GENERator, have demonstrated the power of million-basepair context modeling, they often exhibit a "representational gap" when applied to plant systems (Wu et al., 2026; Lin et al., 2025; Boshar et al., 2025; Brixi et al., 2026; Avsec et al., 2026). The field of plant genomic foundation models has also expanded rapidly, building upon the foundations laid by natural language processing (Mendoza-Revilla et al., 2024; Zhai et al., 2025; Terrail et al., 2026). Early models such as GPN pioneered the use of masked language modeling (MLM) on Brassicales genomes to predict functional constraints in *Arabidopsis thaliana* (Benegas et al., 2023).This was followed by the Agronomic Nucleotide Transformer (AgroNT), a 1-billion-parameter model trained on 48 edible plant genomes (Mendoza-Revilla et al., 2024). AgroNT introduced the Plants Genomic Benchmark (PGB) and showed that large language models could obtain state-of-the-art results for regulatory annotations and gene expression prediction, although its use of non-overlapping 6-mer tokens limited its performance on fine-scale tasks. PlantCaduceus 2 (PlantCAD2) utilizes the Mamba2 backbone to scale linearly with sequence length, supporting contexts up to 8,192 base pairs while maintaining single-nucleotide resolution (Zhai et al., 2025). Simultaneously, the Botanic0 family of models demonstrated that scaling model capacity on plant-only corpora leads to consistent improvements in predictive power (Terrail et al., 2026). It should be noted that substantial differences exist among models in terms of architecture design, data sources, species coverage, and context length. As a result, their performance is often difficult to compare directly, and different strategies mainly reflect trade-offs among computational cost, representational capacity, and task adaptability. PDLLMs (Liu *et al*., 2025) have explored diverse tokenization strategies and model architectures during pre-training. As a result, these models often exhibit different strengths across downstream tasks, with predictive performance varying depending on the specific application.

Generalist models prioritize cross-species coverage but often lack the intra-species depth required to capture the high degree of structural variation, transposable element (TE) turnover, and population-specific regulatory patterns that characterize crop genomes. Existing plant models like AgroNT, PlantCAD2, Botanic0 have made significant strides by training on diverse species sets. Nevertheless, they are often limited by either tokenization strategies that sacrifice single-nucleotide resolution or architectures that struggle to further extending context modeling to the megabase scale and integrating the high-resolution pangenomic variation of specific crop populations (Mendoza-Revilla et al., 2024; Zhai et al., 2025; Terrail et al., 2026). The objective of this study is to construct a rice-centric foundation model that internalizes the complex pangenomic structure of rice. By training on a collection of cultivated and wild rice genomes, a model, OneGenome-Rice (OGR), which captures evolutionary conservation and regulatory logic at a resolution previously unattainable by generalist models, was developed. The emerging of OGR represents a paradigm shift in crop genomics, transitioning from single genome or pan-genomes to crop-centric population-scale genomic foundation models.

## RESULTS

### Model architecture and training

OGR is a generative genomic foundation model designed to process DNA sequences up to 1 million base (Mb) pairs in length. The model features 1.25 billion total parameters and 1 Mb input DNA sequence, utilizing a Mixture of Experts (MoE) architecture that allows for high representational capacity while maintaining computational efficiency during inference (Figure 1, middle panel). OGR is architected as a 12-layer MoE model built upon the transformer paradigm, a design decision substantiated by the well-documented intrinsic performance advantages of MoE architectures over dense counterparts under equivalent computational budgets. The model commences with a token embedding layer that maps discrete nucleotide tokens to continuous vector representations, followed by three strategically interleaved root mean square normalization (RMSNorm) layers to promote training stability. Positional encoding is implemented via Rotary Positional Embedding (RoPE) configured with an elevated base frequency of 50,000,000, endowing the model with the capacity to accommodate context lengths of up to one million tokens; this is achieved by applying rotary transformations directly to query and key vectors during the attention computation, thereby circumventing the need for explicit positional embeddings. The attention mechanism employs grouped-query attention (GQA), wherein 16 attention heads share a reduced set of 8 key-value groups, striking an optimal balance between computational efficiency and representational expressivity. The MoE module comprises a routing network and eight parallel expert subnetworks, each utilizing Swish-Gated Linear Unit (SwiGLU) activations to augment nonlinear expressive capacity. For each input token, the router selectively activates two experts in a content-dependent manner, enabling adaptive distribution of computational resources across heterogeneous genomic contexts. Ultimately, a linear projection layer maps the terminal hidden state to logits spanning the vocabulary space, and a subsequent softmax transformation yields a probability distribution over candidate tokens in accordance with the next-token prediction objective. This architectural framework is inherently flexible, thereby facilitating seamless adaptation to a diverse spectrum of downstream genomic applications. OGR was pre-trained on a curated corpus of 422 rice genomes (Supplementary Table 1), representing a diverse phylogentic group, which includes both modern high-yielding varieties and wild ancestral lines. The architectural details of OGR model and pre-training strategies were summarized at Table 1 (more details see Supplementary Table 2).

**Figure 1.**
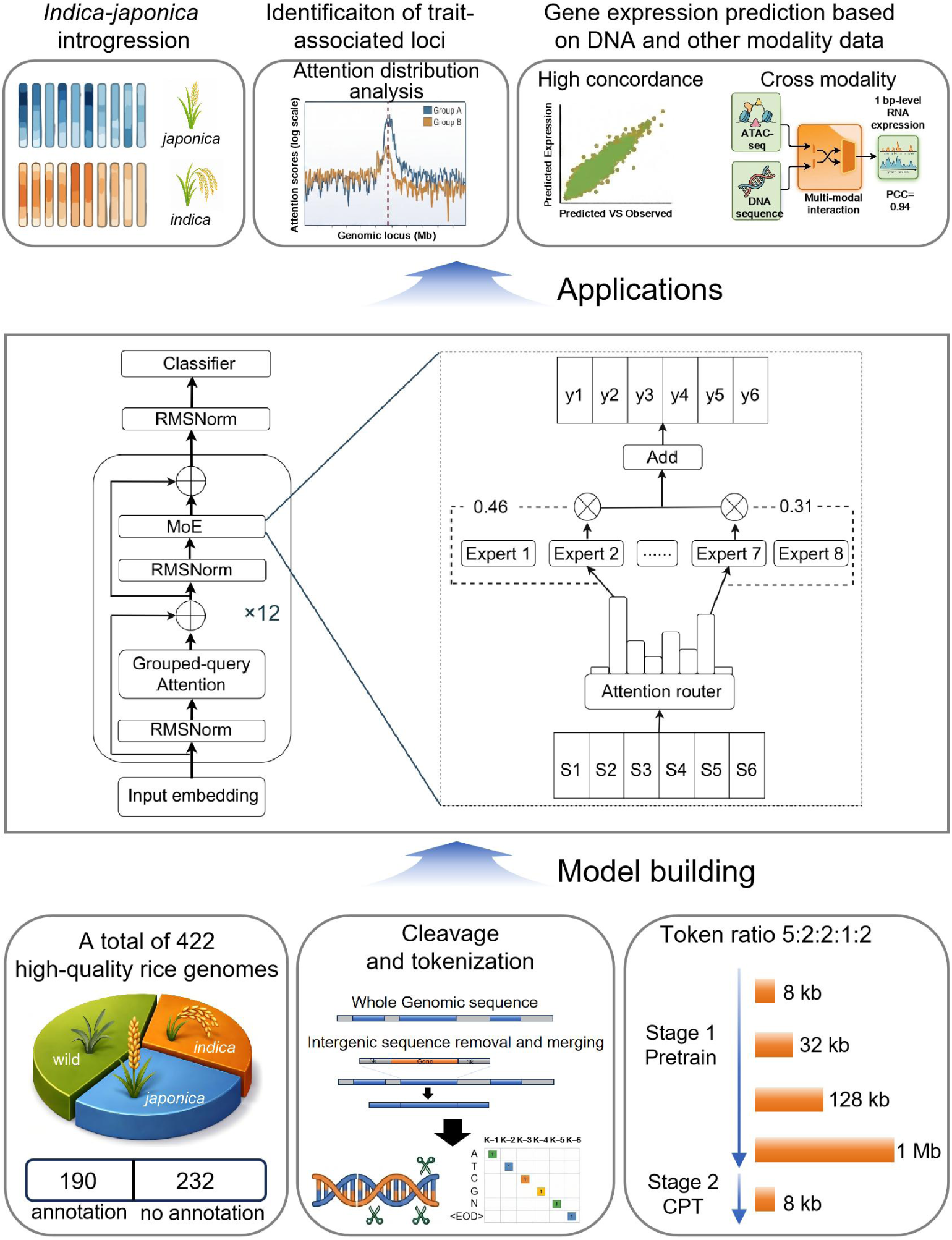
Overview of the OneGenome-Rice (OGR) framework. Conceptual illustration of OneGenome-Rice, a rice genome foundation model trained on population-scale genomic data and applied to diverse downstream tasks, including *indica-japonica* introgression, functional site discovery, and gene expression prediction. Composition of the pretraining dataset, comprising 422 rice genomes spanning *indica*, *japonica*, and wild populations. Model architecture based on a transformer backbone, incorporating tokenized DNA input, positional encoding, and long-range attention mechanisms trained using autoregressive language modeling. CPT: Continuous Pre-Training; MoE: Mixture of Experts.

**Table 1.** Architectural details of OneGenome-Rice (OGR) model.

|  |  |
| --- | --- |
| Architecture | MoE |
| Number of total parameters | 1.25 B |
| Number of activated parameters | 0.33 B |
| Number of layers | 12 |
| Attention hidden dimension | 1024 |
| MoE hidden dimension (per expert) | 4096 |
| Number of attention heads | 16 |
| Number of experts | 8 |
| Selected experts per token | 2 |
| Vocabulary size | 128 (padded) |
| Context length | Up to 1 Mb |
| Attention mechanism | GQA & Flash Attention |
| Activation function | SwiGLU |
| Trained tokens | 594 B |

### Benchmark suite and evaluation framework

To provide a unified assessment of OGR, we used a suite of 26 benchmark datasets across five distinct task categories and additionally incorporated five evaluation tasks from the AgroNT (Mendoza-Revilla et al., 2024). These tasks span multiple levels of genomic function prediction, ranging from local sequence feature identification to the inference of complex population history. We first selected a subset of rice datasets from the AgroNT as evaluation tasks, covering representative functional prediction tasks, including chromatin accessibility, polyadenylation site prediction, and tissue-specific gene expression prediction. Given the large scale of the original benchmark, we randomly sampled 100,000 sequences for the chromatin accessibility task to construct the evaluation dataset, thereby improving computational efficiency while maintaining representative performance assessment (Supplementary Table S3).

To further expand the evaluation scope, we constructed a rice-centered multi-level benchmark framework by systematically designing tasks along two dimensions: sequence length and biological function. Specifically, the benchmark comprises tasks with varying sequence lengths: short-sequence tasks ≤ 1,000 bp (1 kb), long-sequence tasks ≤ 8,000 bp (8 kb), and single-nucleotide tasks (Supplementary Table S3). Overall, these tasks span diverse genomic function prediction scenarios, which can be broadly categorized into epigenetic regulation-related tasks (e.g., chromatin accessibility and histone modifications), cis-regulatory element identification (e.g., enhancers), transcriptional and post-transcriptional regulation (e.g., splice sites and small RNAs), as well as functional element and variant effect prediction. This design enables a comprehensive evaluation of model capabilities across multiple biological and regulatory levels.

In addition, we designed two categories of tasks related to rice evolution and varietal differentiation: selective sweep region identification and rice varieties classification (Supplementary Table S3). The selective sweep region identification task aims to identify genomic regions under evolutionary selection, thereby evaluating the model’s ability to capture population genetic and evolutionary signals. The variety classification task focuses on distinguishing among *japonica* rice, *indica* rice, and wild rice, enabling assessment of the model’s ability to discriminate fine-scale genetic variation among rice population. To systematically investigate the model’s ability to capture long-range dependencies, we introduced multi-scale input sequences for both tasks. Specifically, the selective sweep region identification task employed sequence lengths of 8 kb, 32 kb, and 100 kb, whereas the variety classification task adopted sequence lengths of 8 kb, 32 kb, and 128 kb, enabling comprehensive evaluation across different contextual ranges.

To ensure fair and consistent comparisons across models, sequence embeddings were first extracted using each pretrained model and subsequently used as input to a shared multi-layer perceptron (MLP) classifier or regressor with an identical architecture across all evaluation tasks. Classification performance was evaluated using the Area Under the ROC Curve (AUC), whereas regression performance was measured using the Pearson correlation coefficient.

### Benchmark performance

To systematic evaluate the performance of OGR, we conducted comprehensive comparisons across 26 benchmark datasets/tasks against several representative genomic foundation models, including AgroNT, PlantCAD2, Botanic0, and GENERator-v2 (Table 2). Across the 26 benchmark categories, OGR achieved first- or second-place performance in 16 tasks, demonstrating strong overall capability and robust generalization across diverse genomic prediction scenarios. Notably, OGR exhibited strong performance in multiple regulatory and functional prediction tasks, including chromatin accessibility, histone modification, small RNA prediction, enhancer strength prediction, sweep region identification, and rice variety classification. These results indicate that OGR effectively captures genomic regulatory signals and functional patterns across multiple biological and genomic scales. In contrast, OGR exhibited relatively weaker performance in several fine-grained sequence-level prediction tasks, such as splice site identification, polyadenylation site prediction, and coding sequence (CDS) region prediction, suggesting limitations in modeling highly localized nucleotide-level features. This performance pattern may be partially attributed to the design of the pretraining data and training strategy. During pretraining, OGR was exposed not only to gene-centered sequences with flanking regions but also to large-scale genome-wide sliced sequences, as well as long-context inputs ranging from 32 kb to 1 Mb. Such training strategy likely biases the model toward capturing long-range genomic dependencies and distributed contextual representation, while potentially reducing sensitivity to fine-scale local sequence motifs. This may explain the relatively weaker performance observed in tasks that rely heavily on localized sequence signals, such as splice site, CDS, and polyadenylation site prediction. Conversely, the long-context pretraining strategy appears to provide substantial advantages for tasks that depend on broader genomic context and population-scale variation patterns. In particular, OGR achieved strong performance in sweep region identification and rice variety classification, both of which require integration of long-range genomic dependencies and evolutionary signals distributed across extended genomic regions.

**Table 2.**
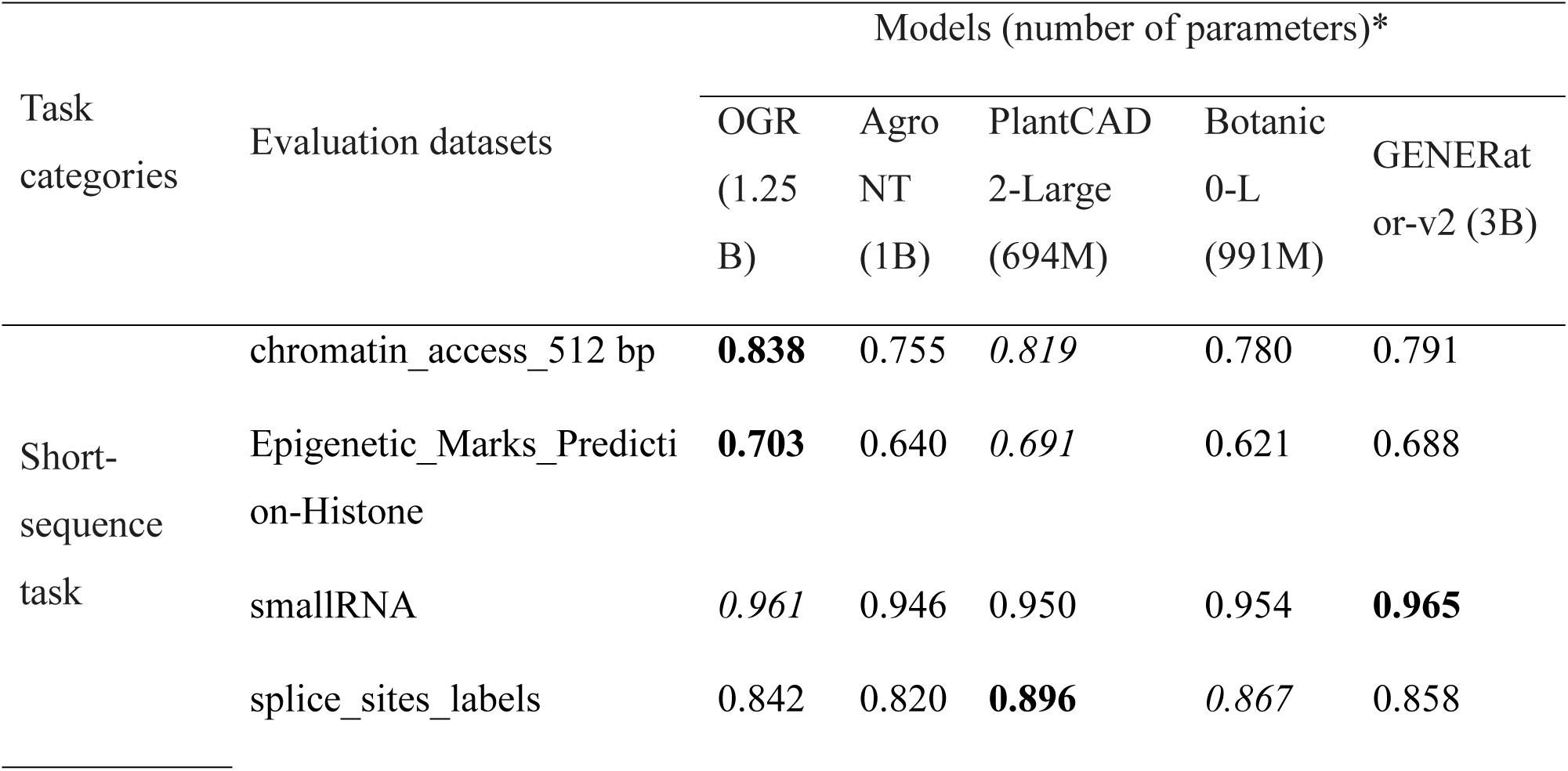
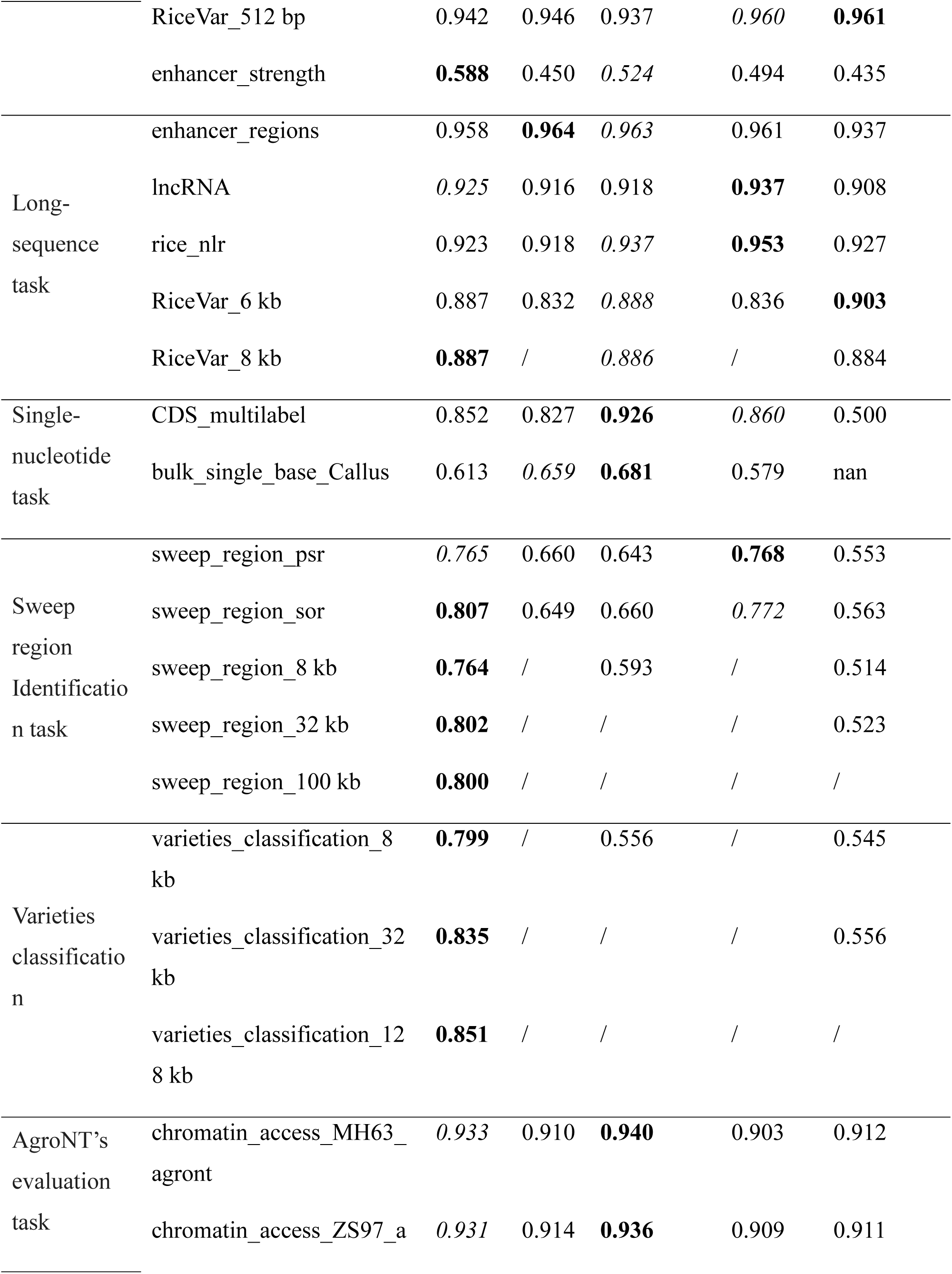
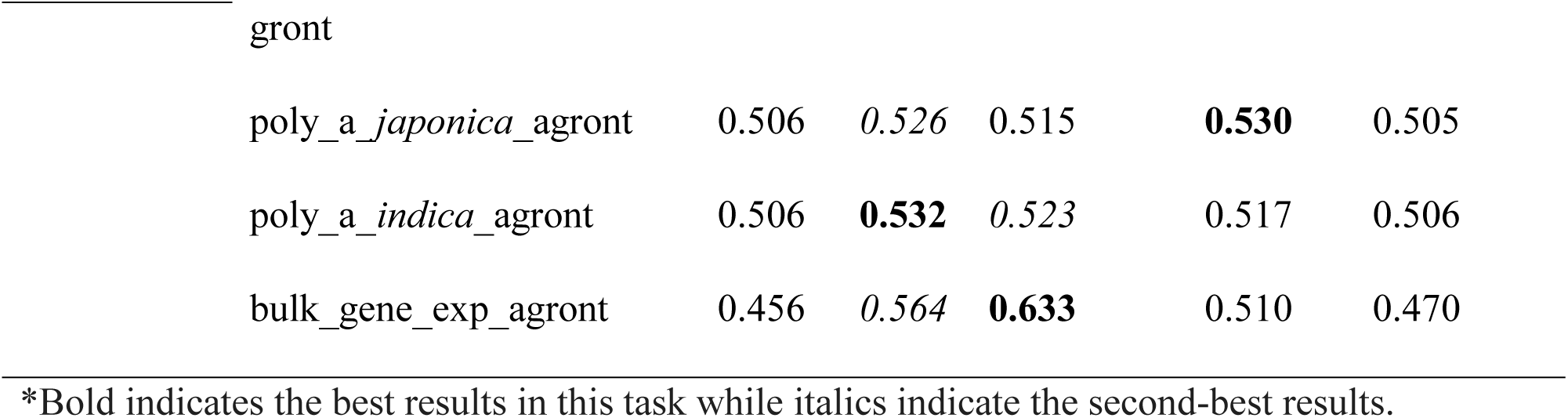
Performance comparison of models on rice benchmark tasks.

Overall, OGR demonstrates strong and balanced performance across diverse genomic prediction tasks, with particular strengths in regulatory sequence modeling and long-context genomic representation learning. These findings suggest that OGR is especially well-suited for applications requiring large-scale genomic context integration, such as population genomics, variety classification and functional genome analysis, while further improvements may be needed for fine-grained nucleotide-resolution prediction tasks.

### Application scenario 1: identification of *indica-japonica* introgression

Modern rice breeders are increasingly utilizing *indica-japonica* heterosis by combining inter-subspecific hybridization and backcrossing to integrate favorable alleles associated with yield, taste quality and adoption (Cui et al., 2022; Sun et al., 2012). Accurate identification of genome-wide introgression enables the dissection of successful subspecific patterns in breeding history, thereby guiding design breeding. However, current phylogenetic methods and SNP-based population comparison analysis are often dependent on reference genomes. They are not readily scalable to newly introduced samples without reconstructing population comparison frameworks, and may be biased in genomic regions with low sequence diversity. So we proposed an alignment- and variant-free framework based on OGR that operates directly on genome assemblies for fine-scale and robust detection of *indica*-*japonica* introgression.

To enable genome-wide identification of subspecies introgression, we constructed a prediction framework based on the OGR foundation model and fine-tuned it in an end-to-end manner for subspecies-origin inference (Figure 2A; details see Methods). We focused on landrace accessions because they better preserve representative subspecies genetic diversity while exhibiting relatively limited confounding from modern breeding-driven admixture. Training samples were selected from the 3K Rice Genome Project dataset (3KRGP) (Wang et al., 2018), including accessions annotated as Chinese landraces, together with a small subset of Japanese landraces and cultivated varieties. Finally, 25 accessions were selected from each of the *indica* and temperate *japonica* subpopulations (Supplementary Table S4). The selected samples were further divided into training, validation, and test sets at a ratio of 3:1:1 for model training and evaluation.

**Figure 2.**
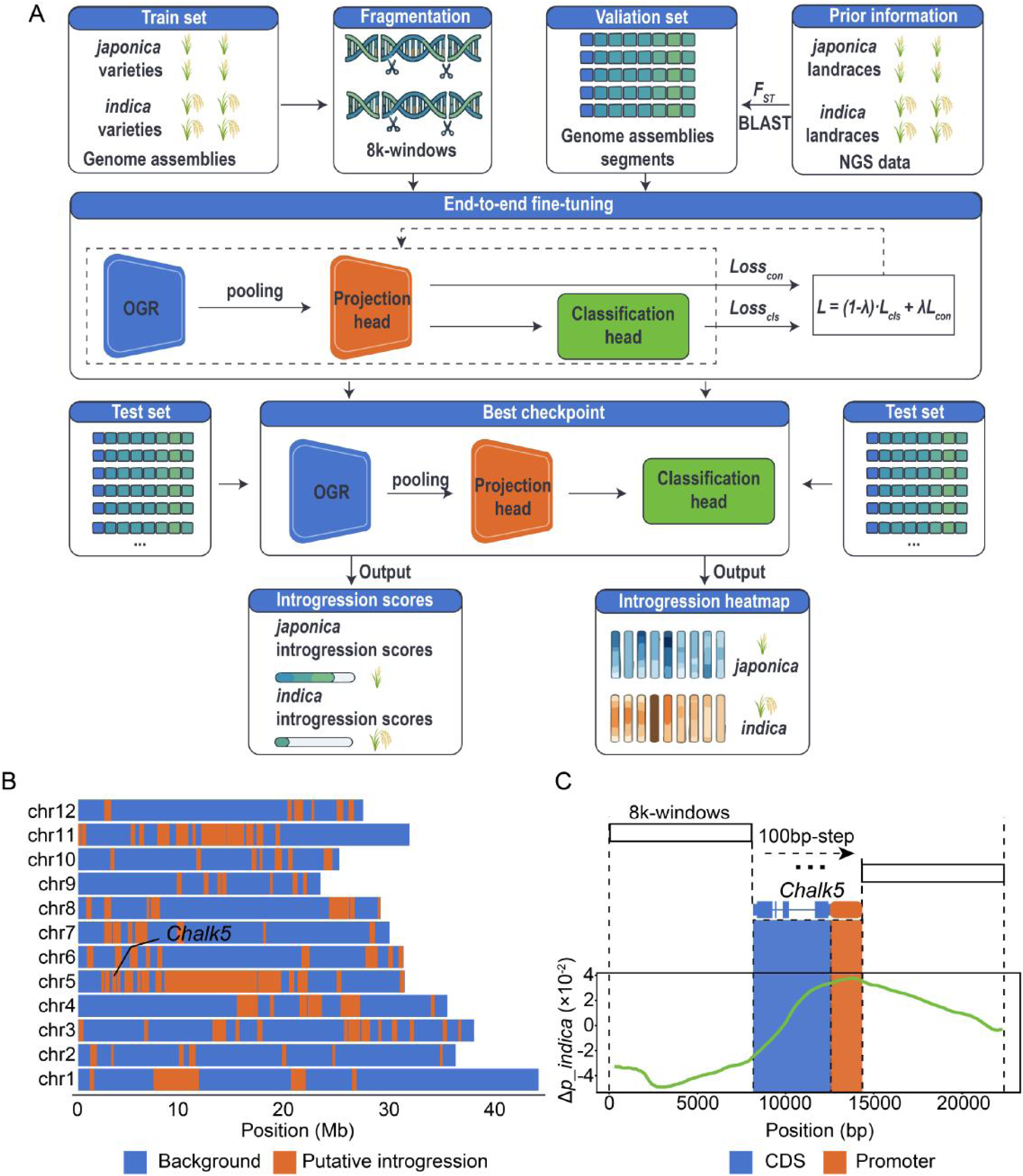
Identification of genome-wide introgression based on OGR. (A) End-to-end fine-tuning framework based on the OGR foundation model for subspecies introgression prediction. Genomic sequences from representative *japonica* and *indica* varieties were segmented into 8 kb sequence windows using a sliding step of 7 kb to construct the training dataset. Highly differentiated genomic regions were identified through *F_ST_* analysis using curated accessions from the 3KRGP accessions, and subsequently projected onto genomic sequences via BLAST to generate high-confidence validation and test datasets. Sequence embeddings extracted by the OGR encoder were subjected to masked mean pooling and then passed into a projection head and a classification head. The model was optimized using a joint objective combining classification loss and contrastive loss. Model checkpoints were selected based on validation performance to ensure robust generalization on the independent test set. The resulting framework generates probabilistic introgression scores for each genomic window, which can be visualized as genome-wide introgression heatmaps. (B) Genome-wide distribution of putative *indica* introgression regions in the elite *japonica* cultivar YF47. *Indica* ancestry inferred by the fine-tuned OGR framework was regionally aggregated, and windows exceeding the confidence threshold (*ε* = 0.5595) were defined as putative introgression regions (orange) against genomic background (blue). The locus surrounding *Chalk5* is labeled. (C) Fine-scale analysis of the *Chalk5* introgression signal. Local ancestry contribution was evaluated using a sliding-window perturbation strategy, and Δ*p_indica* represents the change in inferred *indica* ancestry associated with each 100-bp interval. Gene structure is shown with coding sequence (CDS, blue) and promoter region (orange).

The validation and test datasets were constructed from highly differentiated genomic regions identified through *F_ST_* analysis using curated accessions from the 3KRGP dataset. These differentiated regions were subsequently projected onto the assembled genomes through a BLAST-based alignment procedure, and the top-ranked matching segments were retained to generate high-confidence evaluation datasets enriched for subspecies-divergent signals. Because these regions exhibit strong *indica*-*japonica* differentiation, introgression signals are generally more distinguishable under this evaluation setting. A probability threshold of *ε* = 0.5 was used for evaluation. Using the checkpoint with the best validation performance, the model achieved an accuracy of 78.95% and an AUC of 86.44% on the test set.

We further applied the refined framework to investigate *indica* introgression in the elite *japonica* cultivar Yanfeng 47 (YF47), a representative breeding line shaped by historical inter-subspecific breeding. Rather than adopting an empirically selected cutoff, we estimated the introgression threshold based on the ancestry probability distribution of 8-kb fragments derived from the training set of pure *japonica* landraces. Specifically, the 95th percentile (Q95) of the *indica* introgression probability distribution was selected as the confidence threshold (*ε* = 0.5595), thereby reducing ambiguous predictions arising from genomic regions commonly shared between subspecies. Considering that not every 8-kb fragment contains informative subspecies-differentiating signals, we further quantified local *indica* introgression using a regional aggregation strategy. *Indica* ancestry probabilities were aggregated within 256-kb windows using a 64-kb sliding step, and the average probability of the top 10 highest-scoring 8-kb fragments within each window was used to represent local introgression intensity. Windows exceeding the Q95 threshold were subsequently defined as putative *indica* introgression regions.

Based on this strategy, we identified multiple putative indica introgression regions across the YF47 genome (Figure 2B). The regional aggregation framework improved ancestry signal robustness and facilitated the detection of chromosomal regions potentially reflecting historical introgression and recombination events during breeding. These results provide a genome-wide view of *indica* genomic components within this elite *japonica* background and demonstrate the utility of OGR for introgression identification in breeding materials. Among the detected regions, a prominent introgression signal was identified surrounding the *Chalk5*, consistent with previous observations (Xu et al., 2023).

To further resolve the ancestry contribution at *Chalk5* locus, we performed a fine-scale perturbation scan integrating the original 8-kb ancestry framework with a 100-bp stepwise analysis (Figure 2C). Specifically, the scanning window was progressively shifted from a configuration in which the right boundary aligned with the 3′-untranslated region (3′-UTR) toward the upstream region until the left boundary extended beyond the outer boundary of the 2-kb promoter region. For each 100-bp interval, we quantified its contribution to *indica* introgression inference by calculating Δ*p_indica*, defined as the difference between the average *indica* probability of windows carrying the focal 100-bp segment and that of windows lacking the segment. Positive Δ*p_indica* values therefore indicate sequence intervals that increase inferred *indica* probability. We observed that inferred *indica* ancestry gradually increased as the scanning window entered the *Chalk5-*CDS region, with the strongest contribution localized within the promoter region. This pattern suggests that the *indica*-derived haplotype surrounding *Chalk5* may primarily reside within regulatory sequences rather than coding regions, potentially influencing gene expression through cis-regulatory variation. Overall, these results demonstrate that OGR not only recovers genome-wide ancestry structure through end-to-end fine-tuning, but also enables fine-scale dissection of functionally relevant introgression events. By integrating confidence thresholds, regional aggregation, and locus-level perturbation scanning, the framework provides a practical strategy for identifying breeding-relevant introgression signals in breeding materials.

### Application scenario 2: trait-associated loci finding

The utilization of genomic variation to enhance phenotypic performance is a pivotal objective in modern crop breeding. DNA language model-based approaches for identification of causal variants offer the potential advantages in scenarios where association-based methods encounter challenges due to limited population size, small-effect loci, or low-frequency variants. Inspired by ATLAS (Attention-based Locus Analysis System), a framework developed for human populations that enables population-level discovery through attention patterns from genomic language models (Liu et al., 2026), we extended this concept to rice, and developed an OGR-driven pipeline for genotype-to-phenotype analysis in rice populations.

To validate the pipeline’s ability to identify genes or mutations associated with quality-related traits, we first applied it to glutinousness a trait with a well-characterized genetic basis. In this analysis, we initiated the process by reconstructing sample-specific DNA sequences. This was achieved by integrating variant calls from VCF files with the reference genome. Subsequently, attention scores were then computed separately for glutinous and non-glutinous samples. These scores were then subjected to reference-coordinate normalization and differential analysis at both the site and block levels. Preliminary examination revealed notable differences in attention score distributions across genomic positions between the two phenotypic groups (Figure 3A). To identify candidate regions associated with glutinousness, a two-stage prioritization strategy was employed. Firstly, local peak signals were identified by detecting positions with the strongest differential signals across the analyzed genomic region based on the forward and reverse attention-derived profiles. Secondly, following the mapping of these signals to annotated gene intervals, a series of gene-level metrics were calculated for each gene. These metrics were then subjected to further assessment for the purpose of identifying genes that demonstrated consistent signal enrichment.

**Figure 3.**
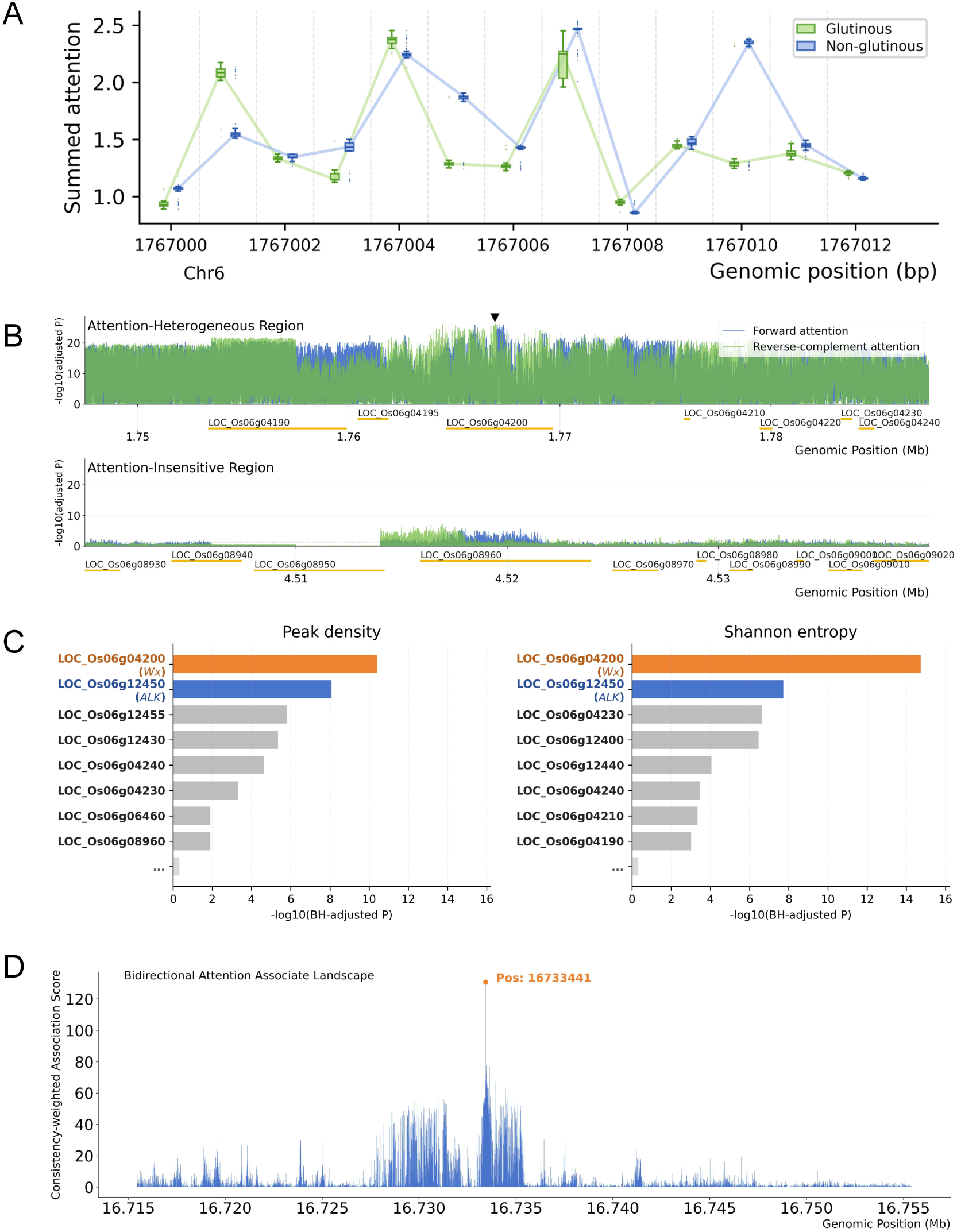
Attention-based study on agronomy trait-associated sites using OGR. (A) Boxplot visualization of summed bidirectional attention scores across a 13-bp window within the major-effect glutinousness locus *Wx* (LOC_Os06g04200). Each genomic position shows the distribution of attention scores in glutinous and non-glutinous rice groups, with connecting lines indicating group medians. (B) Position-resolved differential attention signals in two representative genomic regions in chromosome 6. The upper panel shows an attention-heterogeneous region containing the *Wx* gene (LOC_Os06g04200), whereas the lower panel shows an attention-insensitive comparison region. For each position, forward and reverse-complement attention scores were compared between glutinous and non-glutinous rice groups, and the signal intensity is shown as -log10 (BH-adjusted *P*). The solid black inverted triangle highlights the strongest differential signal in the tested regions, which co-localizes with the *Wx* gene, indicating highly significant group-level attention differences near this major-effect. (C) Gene-level comparison of attention-derived metrics across the selected 160-kb region. For both Peak density and Shannon entropy, *Wx* showed the strongest adjusted-P difference between glutinous and non-glutinous rice groups, while *ALK gene* ranked second. These results suggest that attention-derived regional metrics can help highlight biologically relevant loci associated with rice quality variation. (D) The Manhattan-style landscape displayed illustrates the bidirectional attention association profile of a 40 kb region on chromosome 3 (16,715,000-16,755,000 bp) containing the *GS3* gene, a grain-size regulator. In this region, CALM finds a strikingly sharp and dominant association peak precisely at position 16,733,441, which corresponds exactly to the known QTN underlying the GS3 gene.

As the attention-based outputs represent ranked prioritization results rather than threshold-based association calls, no fixed significance threshold or predetermined number of discovered loci was imposed. Instead, the objective was to evaluate whether the top-ranked intervals and genes captured known biological drivers and highlighting additional candidates of potential interest. At the site level, the most statistically significant differential peak between glutinous and non-glutinous samples was mapped to the *Wx* region (Figure 3B). This outcome aligns with the established role of *Wx*, which encodes granule-bound starch synthase 1 and functions as a major-effect gene regulating amylose synthesis and glutinous endosperm formation (Wang et al., 1995). Furthermore, genome-wide association study (GWAS) performed on the same set of glutinous and non-glutinous accessions also identified the strongest association signal at the *Wx* locus, thereby supporting concordance between the attention-based prioritization and conventional association mapping. At the gene level, multiple interval-based metrics were examined after aggregation of the local attention signals within annotated gene regions. Among the metrics examined, peak density and shannon entropy proved to be the most informative. Both metrics placed *Wx* at the top of the candidate list and highlighted *ALK* as a significant secondary signal (Figure 3C). *ALK* is known to encode soluble starch synthase 2-3 and it has been demonstrated to participate in starch biosynthesis and starch quality regulation (Nakamura et al., 2005).

Building on the detection of a quality-related quantitative trait locus, we next consider a broader and more general challenge in crop genetics. Most agronomic traits of interest are quantitatively inherited and controlled by complex genetic architectures with multiple loci of small to moderate effects. Accurately capturing these signals therefore requires analytical frameworks capable of capturing subtle genotype-phenotype associations under a rigorous statistical modeling framework. To meet this need, we extended the ATLAS framework and developed a new attention-based analytical strategy, termed CALM (**C**asuality inference with **A**ttention-based **L**inear regression **M**odel), specifically designed for quantitative trait mapping based on linear modeling and statistical hypothesis testing (see Methods). Furthermore, CALM introduces a bidirectional integration strategy that combines signals from forward- and reverse-strand scans using a geometric mean significance together with a directional consistency penalty term, thereby improving the robustness of association signals.

To evaluate the performance of CALM on experimentally measured phenotypes, we analyzed grain length from 2,099 accessions within the 18K rice population (Wei et al., 2024). Remarkably, CALM identified an extremely sharp association peak surrounding the well-documented grain-size regulator gene *GS3*, with the highest signal precisely corresponding to the causal nucleotide site (QTN) located at position 16,733,441 (Fan et al., 2009) on chromosome 3 (Figure 3D). This suggested that the attention score-based association framework might possess advantages for fine-scale localization. Unlike traditional variant-based association analyses, which are often constrained by local linkage disequilibrium (LD) patterns, CALM operates on base-level attention signals that may be more sensitive to functionally important structural regions. This property may therefore provide the potential for finer-resolution mapping of causal variants in complex crop genomes.

### Application scenario 3: gene expression prediction based on DNA sequence and multi-modal data

Predicting transcript abundance directly from DNA sequence could address one of the central challenges in molecular biology, which is linking genotype to phenotype through regulatory mechanisms. To evaluate this capability in OGR, we fine-tuned the OGR foundation model using paired genomic and transcriptomic data from three rice cultivars (Figure 4A). Genomic sequences were segmented into 32-kb windows using a sliding-window strategy with a 16-kb step size, and the model was trained to predict transcript abundance directly from DNA sequence. To assess the robustness and generalizability of the model across genomic contexts, prediction performance was evaluated at the chromosome level using an independent test cultivar accession (Xiushui134). Across all 12 rice chromosomes, the model consistently achieved high concordance between predicted and observed gene expression profiles, with log-transformed Pearson correlations ranging from 0.91 to 0.95 (Figure 4B). Notably, prediction performance was stable regardless of chromosomal location, suggesting minimal positional bias in the learned sequence representations. As a proof-of-concept for biological interpretability, we examined *TAC1* (*Tiller Angle Control 1*), a well-characterized quantitative trait locus (QTL) controlling tiller angle in rice (Yu et al., 2007). The predicted expression changes generated by the model were consistent with the known functional effect of a reported 3’-splice site mutation (AGGA→ GGGA) in intron 4, which disrupts proper splicing and alters the 3’-UTR structure (Figure 4C), resulting in a dramatic decrease (-92.71%) in the differential expression region (DER) and a corresponding 15.93% down in predicted transcript abundance (Figure 4D), leading to a decreased expression level of *tac1* mRNA. This example illustrates the ability of OGR to capture genotype-driven expression changes with direct phenotypic relevance.

**Figure 4.**
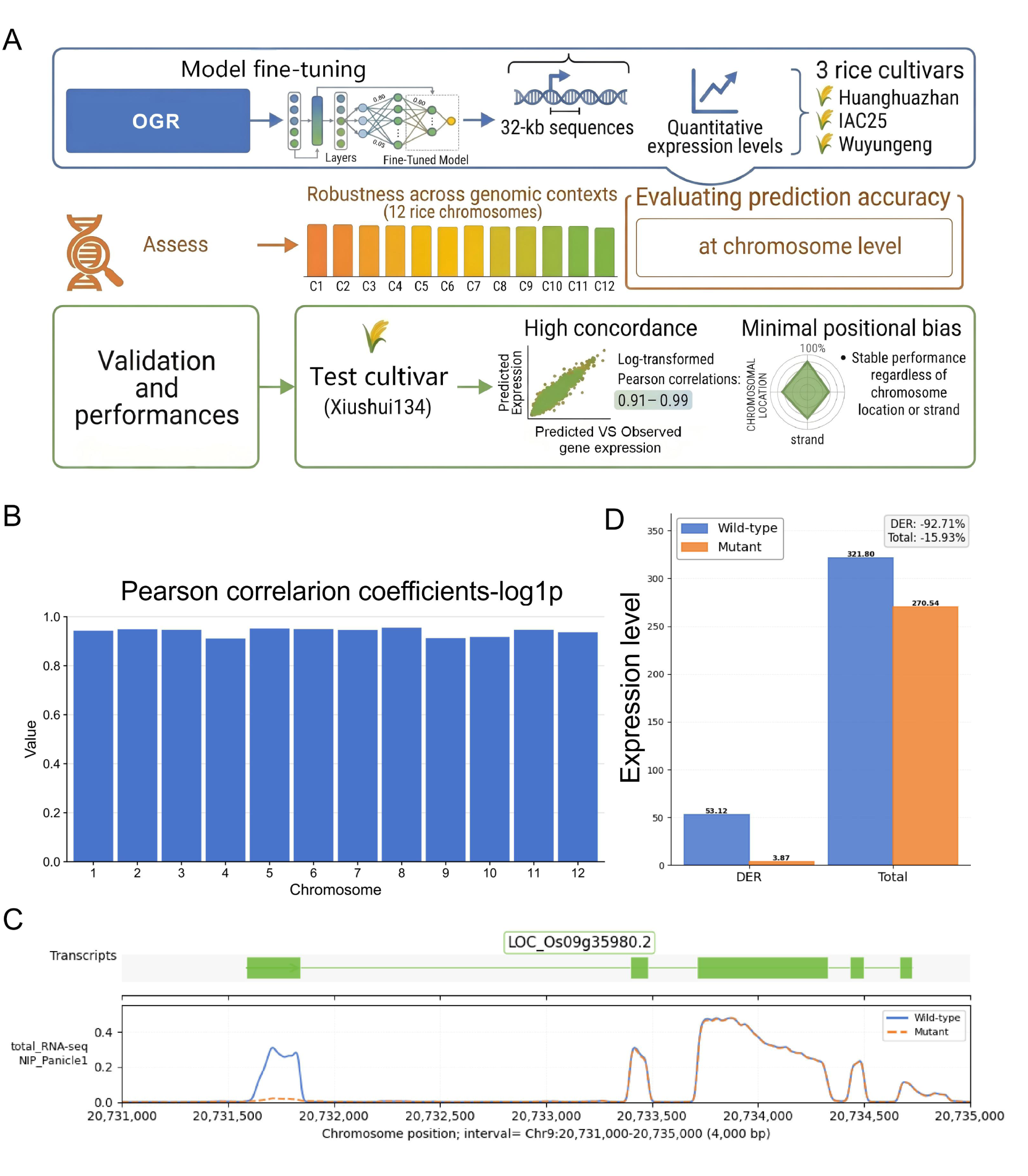
Gene expression prediction of genomic sequence. (A) Overview of the gene expression prediction framework, using fine-tuned OGR to infer gene transcriptional expression levels. (B) Pearson correlation coefficients between predicted and observed expression for the first 8 Mb of each chromosome. (C) The expression prediction of the effect of a nucletide mutation within the *TAC1* gene (LOC_Os09g35980). Predicted expression profiles are compared between the wild-type reference (AGGA; dashed red line) and mutant (GGGA; solid blue line) sequence. A striking divergence occurs around position 20,732,589, where an aberrant signal peak emerges exclusively in the wild type profile. (D) A quantitative comparison of the predicted expression between mutant and wild type. The total expression values are plotted for the differential expression region (DER) (left) and the total region of the gene (right).

Although genomic sequences provide fundamental structural information, the actualization of traits is largely governed by the nuanced interplay between DNA and its regulatory environment. The capability of cross-modal prediction could facilitate preliminary *in silico* perturbation analysis, potentially allowing for the digital simulation of rice responses to environmental stressors or genetic modifications, complementing conventional experimental approaches. To explore this approach, we developed a cross-modal prediction framework consisting of four integrated components (Figure 5A). The pretrained OGR foundation model was employed as the primary DNA encoder, with only the final layer unfrozen during training to utilize its pretrained genomic representations while maintaining computational stability. This was paired with a lightweight transformer tower designed to process chromatin accessibility (ATAC-seq) signals. The DNA and ATAC representations were subsequently integrated via a cross-attention mechanism followed by a four-way gated multilayer perceptron (MLP) fusion layer. The fused representations were then informed a U-Net decoder. To help preserve fine-scale spatial resolution, the decoder utilized residual skip-connections from both the DNA and ATAC encoders, enabling single-base resolution prediction.

**Figure 5.**
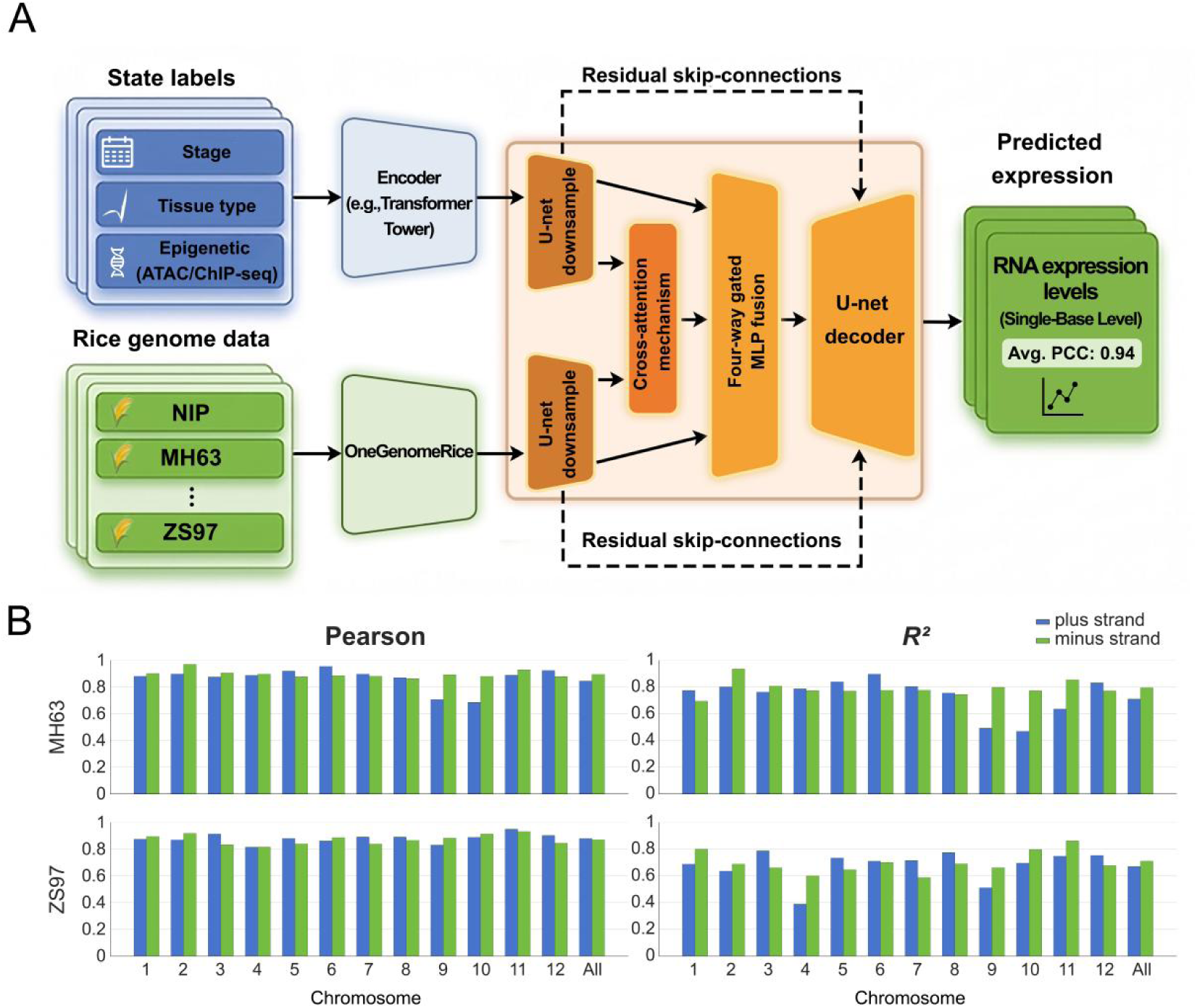
Gene expression prediction based on rice multi-modal data. (A) Schematic of the fine-tuned model architecture for expression prediction. DNA is encoded by the OGR foundation model (final layer trainable; remaining weights fixed). Chromatin accessibility by ATAC-seq is encoded by a lightweight transformer tower. The two representations are downsampled separately and then combined through cross-attention and a four-way gated MLP fusion module, followed by a U-Net upsampler as decoder that outputs single-base resolution predictions. Residual skip connections from the DNA and ATAC encoders feed the decoder to help preserve detail along the sequence. (B) Concordance between single-base-resolution gene expression predictions and observed RNA-seq data in two example varieties (MH63 and ZS97). The paired datasets (DNA and ATAC-seq) from the shoot apical meristem of two varieties at 30 days post-planting were used. Pearson, Pearson correlation coeffeicients. *R^2^*, the coefficient of determination.

The preliminary performance of this architecture was evaluated using a curated dataset containing paired reference genomes, ATAC-seq profiles, and RNA-seq data (Zhu et al, 2024). Experiments were performed using shoot apical meristem samples collected from the MH63 and ZS97 cultivars at 30 days after planting, each with two biological replicates. For each cultivar, the model was trained on one biological replicate and evaluated on the held-out replicate from the same tissue and developmental stage. Under this setting, genome-wide evaluation across both DNA strands showed an average single-base-level Pearson correlation coefficient of 0.880 between predicted and observed RNA expression signals, with chromosome-strand-level values ranging from 0.687 to 0.971 (Figure 5B). This high correlation suggests that the model effectively captures the relative abundance and spatial distribution of the RNA expression spectrum. However, the coefficient of determination (^2^) exhibited greater variability across experiments, with an average value of 0.727 and chromosome-strand-level values ranging from 0.390 to 0.936. This fluctuation highlights the inherent challenge of accurately recovering the absolute scale of RNA expression signals, particularly for data characterized by a high dynamic range (where the maximum-to-minimum ratio exceeds 10^5^). Nevertheless, the overall performance indicates that the model is capable of capturing interactions between chromatin architecture and genomic sequence in the regulation of transcriptional output. Although further validation across tissues, developmental stages, and environmental conditions remains necessary, these initial results suggest that OGR may serve as an effective representation backbone for multi-modal rice genomic modeling and contribute to the continued advancement of AI-driven crop genetics research.

## DISCUSSION

In this study, we present the first rice-specific genome foundation model pre-trained with large-scale cultivated and wild rice genomes. Our results suggest that the large-scale pre-training on a pangenomic dataset (422 accessions) from a single crop species is critical for capturing the species-specific genomic regulatory grammars. Although multi-species genomic foundation models provide broad representations of the central dogma, they frequently lack the sensitivity required to resolve fine-scale population structures, selective sweep detection, and subspecies differentiation in crop genomes. The strong performance of OGR in these tasks confirms that domain-specific scaling, achieved through extensive intra-species variation modeling, is as important as architectural scaling for agricultural genomic applications. The successful deployment of MoE architecture further demonstrates that sparse scaling strategies are effective for modeling the structurally complex and repeat-rich landscapes of cereal genomes. In addition, the integration of RoPE and GQA addresses the long-standing computational challenges of processing megabase-scale genomic sequences at single-nucleotide resolution. Meanwhile, we also provide a standardized rice benchmark suite including 21 new benchmark datasets across five functional categories for future applications of genomic foundation models in rice research and several useful fine-tuned models, including a fine-tuned OGR for alignment-free subspecies introgression mapping, two fine-tuned OGRs for quantitative transcriptomic prediction directly from DNA sequences and multi-modal data, and a new attention-based quantitative trait mapping tool (CALM). Taken together, as a promising foundational computational infrastructure for rice, OGR represents a paradigm shift in crop genomics and precision breeding in future.

Modern rice breeding is rapidly transitioning from empirical selection toward AI-assisted design breeding (Fu et al., 2025), where a central objective is to systematically characterize breeding-relevant genetic resources and thereby enable rational genome-guided crop improvement. In this context, the key challenge is not merely to generate predictive models, but to development AI frameworks capable of transforming genomic information translates into biologically interpretable and breeding-actionable knowledge. Based on OGR, we therefore designed multiple downstream application scenarios that together defined the functional scope of genomic foundation model-driven breeding intelligence.

The indica-japonica introgression analysis demonstrates that OGR can recover population ancestry structure directly from raw genomic sequences without relying on sequence alignment, SNP calling, or predefined population-comparison frameworks. Unlike conventional phylogenetic or marker-based approaches, the OGR-based framework captures subspecies-specific genomic signatures through end-to-end representation learning, enabling robust inference even in regions with low sequence diversity or complex haplotype backgrounds. The observation that introgressed regions in YF47 form extended genomic blocks rather than isolated loci further suggests that the model effectively preserves historical recombination and breeding-derived inheritance patterns. Importantly, the identification of introgression signals surrounding the Chalk5 locus highlights the biological relevance of the learned representations and demonstrates the potential utility of the framework for connecting introgressed haplotypes with agronomically important traits. More broadly, this result indicates that genomic foundation models can provide a scalable and transferable approach for dissecting population structure, tracing breeding history, and identifying functionally relevant introgression events in modern breeding germplasm.

Traditional genetic mapping approaches aim to establish statistical relationships between genotypic and phenotypic variation across populations or segregating populations. However, these methods typically infer trait-associated loci indirectly through linkage disequilibrium between genotyped markers and causal variants. Consequently, many studies resolve broad quantitative trait loci (QTLs) rather than directly identifying functional sequence features responsible for phenotypic variation. In contrast, genomic language model-based approaches such as ATLAS (Liu et al., 2026) and CALM by this study directly utilize genomic sequences as model input and learn contextual representations from nucleotide composition, motif organization, and sequence perturbations induced by genetic variation. This provides a sequence-centered framework for capturing phenotype-relevant genetic signals, complementing conventional association mapping approaches that primarily rely on population-level allele-phenotype correlations. Importantly, because these approaches do not depend on predefined mapping populations or LD structure, they are potentially applicable across diverse genetic contexts, including diversity panels, breeding populations, and mutant populations. Furthermore, by leveraging large-scale pretrained genomic representations and long-range sequence context, genomic foundation models may provide advantages in scenarios where association-based approaches are limited by small population sizes, low-frequency variants small-effect loci or loci with small effect sizes.

Fine-tuning genomic foundation model on paired genomic and transcriptomic datasets further provides important biological insight into how cis-regulatory information is encoded and interpreted within the rice genome. DNA-to-expression prediction established a direct quantitative relationship between sequence variation and transcriptional output, thereby strengthening genotype-phenotype inference and facilitating functional interpretation of noncoding regions. Such models capture regulatory logic shaped by evolution, domestication and breeding selection, while simultaneously offering scalable framework for *in silico* perturbation analysis, functional element discovery, and rational design of gene expression for crop improvement.

More importantly, the transition from descriptive genomics toward predictive biological modeling necessitates computational frameworks capable of bridging static genomic information and dynamic phenotypic expressions. Although genomic sequences offer the structural blueprint of biological systems, phenotyptic manifestation ultimately emerges from interaction between DNA and its regulatory environment. By integrating DNA sequences, chromatin accessibility, and transcriptomic information within a unified architecture, OGR captures the regulatory interactions underlying cellular identity and transcriptional activity. This cross-modal modeling strategy may help distinguish latent genetic potential from realized biological function, thereby offering a more comprehensive representation of how molecular perturbations propagate through regulatory networks to influence complex agronomic traits.

Multi-species genomic foundation models can utilize evolutionary conservation to provide constraints and improve generalization (Karollus et al., 2024). By integrating genomic sequences from diverse species, these models are particularly effective at capturing conserved functional elements and regulatory logic shared across evolutionary lineages. However, their reliance on cross-species consensus signals may dilute species-specific variation and reduce sensitivity to population-level heterogeneity. This limitation becomes especially pronounced when modeling traits shaped by recent evolutionary processes or artificial selection rather than long-term evolutionary conservation. This issue is particularly relevant in rice, where complex population structures, such as *indica-japonica* admixture and extensive artificial selection, play central roles in shaping agronomic traits. Many breeding-relevant alleles are not relatively conserved across individuals due to they have emerged or been enriched during relatively recent breeding programs. Consequently, single-species models is better suited to resolve fine-scale lineage structure, local haplotype variation, and breeding-associated genomic signatures. By focusing specifically on within-species diversity, these models can more accurately capture the genetic architecture characteristic of crop breeding materials. Single-species models also provide advantages for attention-based interpretability analyses. Because the model is trained on a relatively homogeneous sequence distribution, attention-derived importance signals are more likely to reflect phenotype-associated variation within the species itself, rather than broad interspecies divergence. This enables more precise prioritization of candidate loci associated with agronomic traits. In contrast, although multi-species models are powerful for identifying conserved functional regions, they may attenuate subtle yet biologically important variations that are specific to rice populations, particularly variants introduced through recent breeding or introgression events. As a result, signals critical for distinguishing elite breeding germplasm may become partially masked. Collectively, these observations highlight a fundamental trade-off between generalization and sensitivity in genomic foundation modeling. Multi-species models provide robustness through evolutionary conservation, whereas single-species models offer higher resolution for capturing population structure and breeding-relevant variation. Future studies should explore hybrid modeling strategies capable of integrating the complementary strengths of both paradigms. An important open question is how to incorporate cross-species evolutionary information without compromising sensitivity to within-species variation, or how to design architectures that preserve fine-scale population signals while benefiting from conserved functional priors. Addressing these challenges may enable the development of more flexible and biologically grounded AI frameworks for breeding-oriented genomic discovery.

Despite its promising performance, OGR currently has several limitations. OGR is presently restricted to the rice species, and its generalization to other major cereals like wheat or maize has not been fully explored. Cereal genomes vary significantly in genome size, ploidy, and repeat composition, which may require further optimization of the expert-routing strategies and architectural configuration. Additionally, although OGR demonstrates strong performance in transcript abundance prediction, the current framework primarily models basal transcriptional activity and may struggle to capture highly dynamic environmental responses, such as drought, heat stress, or pathogen-induced transcriptional reprogramming, in the absence of explicit environmental conditioning signals. Notably, while long-context pretraining strategy improves performance on tasks involving long-range genomic dependencies, this strategy may reduce sensitivity to highly localized sequence motifs. Consequently, OGR exhibits comparatively weaker performance in fine-scale sequence prediction tasks such as splice site recognition and CDS region identification. Future work may require hybrid architectures capable of jointly modeling both local motif-level features and ultra-long genomic dependencies. Finally, megabase-scale pre-training remain computationally demanding, potentially limiting rapid model iteration and accessibility in resource-constrained research environments. Future versions of OGR will integrate multi-omics data, including single-cell transcriptomics and three-dimensional genome architecture (Hi-C), to refine the model’s understanding of cell-type-specific regulation and long-range spatial dependencies. Ultimately, OGR may serve as a foundational computational framework for AI-driven genome engineering and breeding, enabling rational design of future rice varieties with optimized yield, stress tolerance, and agronomic performance.

## METHODS

### Model architecture implementation

OGR utilizes a Mixture-of-Experts (MoE) architecture derived from the transformer paradigm, comprising 12 layers and optimized for both predictive performance and computational efficiency in genomic sequence modeling. It is well established that the MoE framework confers intrinsic performance advantages that extend beyond mere gains in computational economy, a foundational principle that motivates the design of our genomic model. Prior investigations have demonstrated that, under equivalent computational constraints, MoE-based models achieve lower perplexity and higher accuracy relative to dense baselines (Shazeer et al., 2017). This heightened modeling capacity, attributable to expert specialization and conditional computation over an expansive parameter space, has been consistently validated in large-scale language models (Fedus et al., 2022) as well as in other complex data domains (Shi et al., 2025). Accordingly, we adopted this well-validated architectural strategy without substantive alteration; the empirical performance improvements observed in the OGR model are therefore fully consistent with theoretical expectations and the preponderance of existing evidence.

The model architecture initiates with a token embedding layer that transforms discrete base tokens into continuous vector representations. Subsequently, three strategically positioned root mean square normalization (RMSNorm) (Zhang & Sennrich, 2019) layers are incorporated within the network to enhance training stability by rescaling input activations to a root mean square value of unity, without applying mean recentering. Between the first and second RMSNorm layers, OGR integrates Rotary Positional Embedding (RoPE) (Su et al., 2024) configured with an exceptionally high base frequency of 50,000,000, thereby endowing the model with the capacity to process ultra-long sequences of up to one million tokens. Notably, in lieu of explicit positional embeddings at the input stage, RoPE encodes positional information dynamically during the attention computation by applying rotary transformations directly to the query and key vectors. This design confers precise positional awareness while accommodating extreme context lengths.

Complementing the RoPE mechanism, the model employs a grouped-query attention (GQA) (Ainslie et al., 2023) scheme comprising 16 attention heads that collectively share 8 key-value groups. This configuration achieves an optimal trade-off between computational efficiency and representational expressivity, enabling accurate and efficient processing of extensive genomic sequences. OGR adopts a MoE architecture consisting of a router network alongside eight parallel expert subnetworks. Each expert subnetwork utilizes the Swish-Gated Linear Unit (SwiGLU) (Zhai et al., 2022) activation function in place of conventional alternatives such as the Rectified Linear Unit or Gaussian Error Linear Unit (ReLU/GELU), thereby enhancing both expressive power and training stability. The router dynamically assigns two of the eight experts to each token in a content-dependent manner, facilitating adaptive allocation of computational resources (Figure 1, middle panel). This design supports efficient handling of both structurally simple repetitive regions and functionally complex regulatory elements.

Finally, a linear output layer projects the model’s terminal hidden state onto logits spanning the vocabulary space, where a softmax transformation subsequently yields a probability distribution over candidate tokens, consistent with the next-token prediction objective (Radford & Narasimhan, 2018). A salient advantage of this architectural framework is its inherent flexibility, which readily permits effective adaptation to a diverse array of downstream applications.

### Training procedure

During the pretraining phase, OGR was optimized under a self-supervised learning paradigm, wherein the model was trained to minimize the next-token prediction objective while concurrently learning generalizable genomic sequence representations. The training procedure was executed using the Megatron-LM framework (Shoeybi et al., 2020) distributed across 128 graphics processing units (GPUs), leveraging an advanced five-dimensional parallelism strategy that integrates tensor parallelism, pipeline parallelism, context parallelism, data parallelism, and expert parallelism. The global batch size was set to 1,024, realized through gradient accumulation over micro-batches of size 1. Parameter optimization was performed via the AdamW (Loshchilov & Hutter, 2019) algorithm, employing a distributed sharded implementation to manage optimizer state storage efficiently. The learning rate was scheduled according to a cosine decay trajectory, incorporating a 5% linear warm-up phase and attaining a peak value of 1×10^−4^, with gradient clipping enforced at a threshold of 1.0 and weight decay set to 0.1.

To mitigate the inherent challenge of expert load imbalance within the MoE architecture, an issue exacerbated by the constrained genomic vocabulary comprising only four nucleotide bases. We implemented an expert load-balancing mechanism supplemented by an auxiliary loss (Shazeer et al., 2017) with a coefficient of 1×10^-3^. This regularization strategy averts router collapse and promotes uniform expert utilization across heterogeneous genomic contexts. Additionally, a Z-loss (Zoph et al., 2022) penalty, applied to the router logits with an identical coefficient of 1×10^-3^, serves to suppress numerical instability and ensure smoother optimization dynamics within the MoE components.

To enable modeling of ultra-long sequence contexts spanning up to one million tokens, a multistage progressive training regimen was employed. This strategy entailed three principal technical measures: incremental exposure to training sequences of progressively extended lengths, scheduled learning rate annealing to counteract catastrophic forgetting (Wang et al., 2025), and application of Rotary Positional Embedding (RoPE)-based context window scaling techniques.

Numerical stability and training fidelity were further enhanced through the adoption of mixed-precision training. Specifically, the BF16 data format was utilized for the majority of arithmetic operations, whereas full FP32 precision was strictly preserved for a subset of critical computations, namely: (a) the Softmax activation within the attention mechanism, (b) gradient accumulation and All-Reduce collective communications, and (c) MoE routing operations. Concurrently, reduced-precision matrix multiplication via BF16 was explicitly deactivated for these sensitive components.

The synergistic integration of Grouped-Query Attention (GQA) and Flash Attention (Dao et al., 2022) endows OGR with complementary advantages. GQA furnishes architectural refinements essential for the efficient partitioning of key-value representations, while Flash Attention provides a highly optimized computational kernel for accelerated attention score evaluation. This concerted design establishes a robust foundation for sustaining high-performance pretraining at scale with expansive contextual windows.

Additional system-level optimizations encompassed the utilization of grouped general matrix multiplication (GEMM) operations (Zhai et al., 2023) to facilitate efficient batched expert computation within MoE layers, the deployment of all-to-all token dispatching protocols (Hwang et al., 2023) for MoE inter-expert communication, overlapping of parameter aggregation and gradient reduction steps to minimize communication latency, and implementation of a cyclic data loader provisioned with eight worker processes to ensure uninterrupted streaming of training data throughout large-scale pretraining.

The pretraining of the model was conducted on the National Artificial Intelligence Public Computing Power Open Innovation Platform, specifically utilizing the Nanhu Computing Framework, which comprises a resource pool of over 1,000 NVIDIA A100-80GB GPUs. For the pretraining phase, a subset of 16 computational nodes was allocated, each provisioned with 8 NVIDIA A100-80GB GPUs, yielding a total of 128 GPUs dedicated to the training workload. The entire pretraining procedure was completed over a duration of approximately eight days.

### Dataset construction

The pre-training corpus for OGR was meticulously constructed to bridge the "representational gap" prevalent in existing plant models. We integrated a pangenomic dataset of 422 high-quality rice genomes, all of which were sourced from open-access data in previously published literature (Guo et al., 2025; Kawahara et al., 2013; Qin et al., 2021; Wang et al., 2023; Wei et al., 2024; Xie et al., 2025; Zhang et al., 2022). These genomes were selected from a larger pool through a rigorous quality control pipeline. To ensure high data fidelity, we only retained assemblies anchored to 12 chromosomes that met stringent criteria: a scaffold N50 > 20 Mb (continuity), a BUSCO score > 95% (completeness), and fewer than 200 gaps with a total gap length < 1% of the genome (correctness).

This curated dataset captures the profound genetic diversity within the *Oryza* genus. Cultivated Rice (257 accessions): this subset encompasses diverse *indica* and *japonica* landraces and improved varieties, reflecting the genomic signatures of environmental adaptation and anthropogenic selection over millennia. Wild Rice populations (165 accessions): we included wild relatives, primarily represented by *O. rufipogon*. These populations serve as pivotal reservoirs of beneficial alleles for biotic and abiotic stress resistance traits frequently lost during the domestication bottleneck. Within this dataset of 422 high-quality genomes, 190 are professionally annotated, while the remaining 232 consist of *de novo* assemblies without annotation. This pangenomic approach ensures that OGR transcends the limitations of the traditional Nipponbare reference genome. By incorporating diverse genomic architectures, the model achieves a comprehensive understanding of global structural variations (SVs) and rare SNPs that are often absent in a single reference.

The training data for OGR were encoded directly from genomic sequences. Each genomic sequence was transformed using a one-hot encoding scheme, with a vocabulary comprising the four canonical nucleotides (A, T, C, and G), the ambiguous base N, and specialized tokens such as <EOD> to delineate sequence boundaries. To foster a general-purpose representation of the rice pangenome, the corpus remained strictly "agnostic" during this stage, no cell-type-specific labels, epigenetic features, or functional annotations were introduced. This approach ensures that the model captures the endogenous logic of the genome, remaining unbiased by experimental conditions or specific biological contexts.

The initial pre-training phase processed approximately 490 billion (490B) tokens through a multi-scale curriculum learning framework. The dataset was partitioned into four escalating window sizes 8,192 bp, 32,768 bp, 131,072 bp, and 1,024,000 bp, distributed at an approximate ratio of 5:2:2:1. Different sampling strategies were applied at different scales to balance functional feature learning and large-scale structural modeling. The 8 kb and 32 kb datasets were generated using a hybrid strategy that combined two data sources: gene-centered fragments from 190 annotated genomes, including gene bodies together with 3 kb upstream and downstream flanking regions; and whole-genome sequences from 232 high-quality unannotated genomes. These sequences were segmented into fixed-length samples of 8 kb and 32 kb for training. In contrast, the 128 kb and 1 Mb datasets were constructed exclusively from all 422 high-quality whole-genome sequences. The genomes were tiled into long-context samples to enable the model to capture large-scale pangenomic variation patterns.To enhance strand robustness and directional invariance, reverse-complement augmentation was applied throughout all training stages. The forward-to-reverse strand ratios for the 8 kb, 32 kb, 128 kb, and 1 Mb stages were set to 1:1, 1:3, 2:2, and 3:1, respectively (Supplementary Table S2).

In the subsequent continued pre-training (CPT) stage, an additional 104 B tokens were generated by tiling the whole-genome sequences of the 190 annotated genomes into fixed-length 8 kb windows, with a forward-to-reverse strand ratio of 1:3. The return to 8 kb context was motivated by the prevalence of short-sequence downstream tasks, which benefits Fine-Tuning performance while preserving the long-range capacity acquired during initial pre-training. The 190 annotated genomes were chosen specifically because their whole-genome sequences had not been exposed to the model previously; the initial 8 kb stage had drawn only from gene-centered fragments of these assemblies, leaving their intergenic regions as novel signal for CPT.

### Benchmark datasets

#### Rice benchmark construction

To systematically evaluate the representational capacity and generalization performance of different models in rice functional annotation, we established a comprehensive benchmark encompassing multiple tasks, including gene structure prediction, epigenetic regulation, and variant effect assessment.

#### Splice site recognition dataset

*splice_sites_labels*: Constructed using the MSU7 genome annotation. Positive samples consist of 400 bp sequences spanning 200 bp upstream and downstream of donor or acceptor sites, while negative samples are sequences containing GT/AG dinucleotides that do not correspond to true splice sites.

#### Coding sequence annotation dataset

*CDS_multilabel*: Constructed using IRGSP1.0 annotation. Sequences were segmented using 512 bp windows, with base-level binary labels assigned. Training and validation sets were partitioned by chromosome, with chromosomes 9 and 12 used as the validation set.

#### Chromatin accessibility prediction datasets

*chromatin_access_MH63_agront* and *chromatin_access_ZS97_agront*: Each benchmark dataset was constructed by randomly sampling 100,000 sequences from the AgroNT evaluation tasks.

*chromatin_access_512*: Sequences of 512 bp were extracted from the RiceENCODE database, with chromatin accessibility labels assigned across four tissues: mature leaves, panicles, roots, and young leaves.

#### Gene expression prediction datasets

*bulk_gene_exp_agront* (gene-level expression): 6,000 bp sequences from AgroNT evaluation tasks, with the goal of predicting gene expression levels across different tissues.

*bulk_single_base_Callus* (single-base resolution): 6,000 bp sequences with bigWig coverage used as regression labels, derived from callus tissue, as reported in (Zhu et al., 2024).

#### Polyadenylation site prediction datasets

*poly_a_japonica* and *poly_a_indica*: Datasets for alternative polyadenylation site prediction tasks from the AgroNT evaluation tasks.

#### Variant effect prediction datasets

*RiceVar_512 bp*, *RiceVar_6 kb*, and *RiceVar_8 kb*: These datasets were constructed based on annotations from the RiceVarMap database, integrating predictions from SnpEff, PolyPhen-2, and SIFT. Only variants consistently annotated as either deleterious or benign by all three tools were selected. Sequence contexts of 512 bp, 6,000 bp, and 8,192 bp were used, respectively.

#### Epigenetic modification prediction dataset

*Epigenetic_Marks_Prediction-Histone*: Based on 13 histone modification datasets from RiceENCODE database, 1 kb genomic windows centered on peaks were extracted. Overlapping regions were merged, resulting in a benchmark task where each sequence can carry multiple histone modification labels simultaneously.

#### Enhancer identification and strength prediction datasets

*enhancer_regions*: Rice enhancer training data used in the RicENN model (Gao et al., 2022).

*enhancer_strength*: Data from Sun et al. (2019), with enrichment values used as labels.

#### Selective sweep region identification datasets

*sweep_region_psr*, *sweep_region_sor*, *sweep_region_8 kb*, *sweep_region_32 kb*, *sweep_region_100k*: Selective sweep regions were obtained from (Jing et al., 2023). Sequences corresponding to the putative selective sweep regions (PSR) and single-origin selective sweep regions (SOR) were extracted, and datasets were constructed with four sequence lengths: 6 kb, 8 kb, 32 kb, and 100 kb.

#### NLR gene identification dataset

*rice_nlr*: NLR genes were obtained from Stein et al (Stein et al., 2018).

#### Non-coding RNA identification datasets

*lncRNA*: Sequences were obtained from the PLncDB database.

*smallRNA*: Sequences were obtained from the CSRDB and sRNAanno databases.

#### Variety classification datasets

varieties_classification_8 kb, varieties_classification_32 kb, and varieties_classification_128 kb: Based on three representative rice genomes (NIP, MH63, and GWHERKH), sequences were extracted using sliding windows of 8 kb, 32 kb, and 128 kb. One-hot encoding was used to label *indica*, *japonica*, and wild rice.

### *Indica-japonica* introgression identification analysis

Based on the sequence representation capabilities of the OGR foundation model, this study constructs a prediction framework that maps genomic sequences to subpopulation origin probabilities, thereby enabling the inference of ancestry introgression between *indica* and *japonica*. The overall workflow is described below:

#### Data preparation and processing

For this study, a curated whole-genome rice dataset was constructed from the 3KRGP, comprising 25 *indica* and 25 *japonica* accessions. The selected accessions included Chinese landraces with clear population annotations and a smaller subset of japonica accessions from Japan. Within each subspecies, accessions were divided into training, validation, and test sets at a ratio of 3:1:1, resulting in 15, 5, and 5 accessions respectively (with balanced number across subspecies). All sequence windows derived from the same accession were strictly assigned to the same data subset to prevent data leakage and ensure independence and generalizability.

For the training set, quality control was first applied to remove consecutive ambiguous bases (N) at both chromosomal ends. A sliding window strategy was then used to segment the genomes, with a window size of 8,000 bp and a stride of 7,000 bp, enhancing local sequence continuity and improving the detection of potential introgression boundaries. The resulting sequences were encoded using the same tokenizer as that used for the pre-trained OGR model.

For the validation and test sets, we first identified genomic regions with extremely strong genetic differentiation (*F_ST_* > 0.9) between *indica* and *japonica* based on genome-wide scan results from the 3KRGP, and constructed a reference set from these regions. Subsequently, BLAST was used to align sequences from the validation and test sets against this reference set, retaining only high-confidence matching regions as candidate analysis windows, thereby ensuring that the model focuses on genomic regions with strong *indica*-*japonica* differentiation signals. To standardize input length, fragments longer than 8,000 bp were truncated, while fragments shorter than 8,000 bp were padded by extracting flanking sequences based on their original genomic coordinates. Finally, all processed sequences were encoded using the same tokenizer as the OGR pre-training stage before being fed into the model for evaluation.

In addition to the above test set, a genome-wide analysis dataset was constructed using the elite *japonica* cultivar Yanfeng 47 (YF47). For this genome, consecutive ambiguous nucleotides (N) at both chromosomal termini were first removed. The cleaned genome was then partitioned into non-overlapping sequence windows using a sliding-window strategy with a window size of 8,000 bp and a step size of 8,000 bp. All window sequences were encoded using the same tokenizer as that used in the OGR pre-training stage.

#### Model architecture

This study adopted the pre-trained OGR model (rice 1.25 B, 8 kb context) as the DNA backbone encoder to extract sequence representations from 8,000 bp genomic windows. During the fine-tuning phase, a Low-Rank Adaptation (LoRA) parameter-efficient fine-tuning strategy was employed. Low-rank trainable parameters were applied only to the q_proj and v_proj layers within the Transformer attention layers, with a rank of 16, scaling factor of 32, and dropout rate of 0.1.

For the token-level hidden states output by the backbone network, masked mean pooling was applied to obtain window-level feature representations. A two-layer fully connected projection head was used to map the feature dimension from 1024 to 512 and further project it into a 128-dimensional embedding space, forming a representation suitable for supervised contrastive learning. A classification head was then attached to output logits for the two classes, which were converted into source probabilities [*P_japonica_*, *P_indica_*] via a sigmoid function. The *japonica* label was defined as [1, 0], and the *indica* label as [0, 1].

The model was optimized using a joint loss function combining classification loss and supervised contrastive loss:

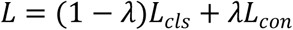

Where *L_cls_* is the BCEWithLogitsLoss classification loss, and *L_con_* is the supervised contrastive loss, which encourages intra-class compactness and increases inter-class separation in the embedding space, thereby improving discriminability in boundary regions between subspecies. *λ* is the loss weight coefficient, set to 0.1 in this study.

#### Training procedure

We trained the model using Distributed Data Parallel (DDP) with NCCL communication backend. FP16 mixed precision was adopted to improve efficiency and reduce memory usage. The model was optimized using AdamW with a learning rate of 1×10⁻⁵ and a cosine decay schedule. Warmup ratio was 0.05, weight decay was 0.01, batch size was 4, and gradient accumulation was 8.

#### Introgression map construction and performance evaluation

For ancestry inference in the test set, a probability threshold *ε* was introduced to evaluate each window, maintaining consistency with the subsequent introgression analysis. Specifically, genomic windows were classified as *japonica*-derived when *P_japonica_* ≥ *ε* and *P_indica_* < *ε*, whereas windows were classified as *indica*-derived when *P_japonica_* < *ε* and *P_indica_* ≥ *ε*. When both probabilities were simultaneously greater than or equal to *ε* or simultaneously less than *ε*, the region was categorized as an uncertain region, indicating ambiguous or low-confidence ancestry inferences.

The discretized ancestry predictions were subsequently mapped back to genomic coordinates to construct discrete label sequences and corresponding probability trajectories at the chromosomal scale, thereby generating a genome-wide ancestry introgression map. Model performance was quantitatively evaluated using the Area Under the ROC Curve (AUC) and Accuracy (ACC).

### Identification of trait-Associated loci

#### Identification and prioritization of glutinousness-associated Loci

To evaluate whether OneGenome-Rice-derived sequence-response profiles could prioritize genomic regions associated with rice quality traits, we applied the proposed pipeline to glutinousness, a binary trait with a well-established genetic basis. Phenotypic information for glutinous and non-glutinous accessions was obtained from the Rice SNP-Seek Database (Mansueto et al., 2017). The corresponding accessions were selected from the 3,000 Rice Genomes Project panel (Wang et al., 2018),, and genome-wide variant files were obtained from RiceVarMap V2.0 (Zhao et al., 2021). Variant calls were retained in a sequence-reconstruction format that preserved SNPs as well as indel/deletion-related variation, allowing accession-specific pseudo-genomic sequences to more closely reflect the observed genomic background. Only accessions with unambiguous glutinousness phenotypes and available genotype data were included, resulting in a final dataset of 100 glutinous and 100 non-glutinous rice accessions.

The genome was partitioned into sliding windows defined in BED format, where each interval specifies the chromosome, start coordinate, and end coordinate of the analyzed genomic block. For each accession and each genomic window, a sample-specific pseudo-sequence of DNA was reconstructed by integrating VCF-derived variants into the reference genome sequence. During this process, an explicit coordinate mapping was maintained between the reconstructed pseudo-sequence and the original reference genome, enabling model-derived scores to be projected back to reference genomic coordinates.

Each reconstructed sequence was processed by OGR in both the forward orientation and the reverse-complement orientation. Model-derived attention or sequence-response scores were extracted separately from the two directions. These scores were then projected back to reference coordinates and reorganized into sample-by-position matrices for each genomic block. Forward-and reverse-direction profiles were retained independently throughout the initial analysis, allowing the consistency of strand-specific signals to be evaluated in subsequent steps.

For each genomic position within a block, sequence-response values were compared between the glutinous and non-glutinous groups. Because glutinousness was treated as a qualitative trait, group differences were assessed using the Mann-Whitney U test. Resulting P values were adjusted for multiple testing using the Benjamini-Hochberg procedure, and the adjusted values were transformed into standardized positional differential signals. This analysis generated forward- and reverse-direction differential profiles across each genomic block, representing the extent to which local sequence-response patterns differed between the two phenotypic groups.

Candidate regions were first identified using a local turning-point interval detection strategy. In brief, the forward- and reverse-direction differential profiles were scanned independently to detect positions showing abrupt local changes in signal intensity. Candidate turning points were defined by local shifts in mean signal together with asymmetry in the surrounding signal distribution. Forward- and reverse-direction candidate points were then paired according to predefined distance and directional-consistency constraints. Paired signals were used to define high-confidence candidate intervals, which were subsequently ranked according to their combined local differential intensity and cross-direction consistency.

To further prioritize candidate genes, positional differential signals were aggregated within annotated gene intervals, allowing attention-derived signals to be summarized at the gene level using reference genome annotations. For each gene, differential signals within the gene body and optional flanking regions were summarized using a panel of statistical features describing signal magnitude, dispersion, distributional shape, peak enrichment, and signal complexity. These included metrics such as central tendency, upper quantiles, peak-related measurements, peak density, and Shannon entropy. The resulting gene-level feature matrix was used to rank candidate genes whose annotated regions showed consistent enrichment of differential sequence-response signals between glutinous and non-glutinous accessions.

Because the attention-derived outputs were designed for candidate prioritization rather than conventional threshold-based association testing, no fixed genome-wide significance threshold or predetermined number of loci was imposed. Instead, the analysis focused on whether the highest-ranked intervals and genes recovered known biological regulators of glutinousness and starch quality, while also identifying additional candidate regions for downstream interpretation.

#### Casuality inference with attention-based linear regression model (CALM)

Building upon the ability of the OGR foundation model to capture functional motif, genomic structure and contextual sequence dependencies, we developed an attention score-based association mapping framework termed as CALM (casuality inference with attention-based linear regression model). Briefly, reconstructive individualized genomic sequences based on VCF files were processed using the pretrained OGR model, and the resulting position-wise attention scores were used for downstream analysis. Unlike traditional GWAS, which uses discrete genotypes as explanatory variables, CALM directly employs attention scores as quantitative factors in a linear regression framework. Specifically, for each genomic position *i*, the following model was fitted:

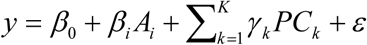

where *y* denotes the phenotypic value, *A_i_* represents the attention score at genomic position *i*, *β_i_* is the estimated effect size, and *^ε^* represents the residual error. Here, principal component analysis was performed directly within the attention-derived feature space to represent population structure and potential confounding effects. *PC_k_* corresponds to the *k*-th principal component calculated from the genome-wide attention matrix. Next, for each genomic position, statistical significance was evaluated by testing the corresponding regression coefficient using a *t*-test, and the resulting *p*-values were used to quantify association signals with the phenotype.

To suppress fake orientation-dependent signals, CALM incorporated a bidirectional consistency analysis. Both the forward genomic sequence and its reverse-complement counterpart were independently analyzed using the same pipeline, yielding forward-direction and reverse-direction association statistics for each genomic position. The two directional signals were integrated using a consistency-constrained framework, in which the final association significance was defined as:

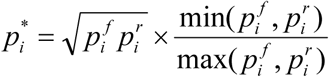

where 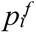 and 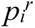 denote the association *p* values calculated from the forward and reverse-complement *t*-test at genomic position *i*, respectively, and *p_i_\** represents the final consistency-constrained integrated significance score. Through this framework, loci showing strong significance in one sequence orientation but weak support in the opposite direction receive greater penalization due to reduced directional consistency. In addition, genomic positions exhibiting inconsistent effect directions between forward and reverse analyses were excluded as non-robust association signals, thereby reducing potential false-positive associations arising from strand-dependent artifacts or local sequence bias. Overall, by performing association analysis directly on contextualized sequence representations rather than isolated genotype states, CALM is less constrained by conventional linkage disequilibrium structure and can improve the resolution of candidate locus localization, thereby providing enhanced fine-mapping capability for identifying potentially functional genomic regions.

### Gene expression prediction

The RNA-seq data from 18K-rice population study were used (Wei et al., 2024). Only leaf tissue data collected 2 to 3 days before heading date were used for analysis in this study. The reference genomes for the parental lines were downloaded from the data repository provided in the original publication. To prevent data leakage, training and test datasets were separated by accession prior to model training.

#### Data preparation

RNA-seq data from three rice cultivars (Huanghuazhan, IAC25, Wuyungeng) at the grain-filling stage were used for model training, wherase a fourth cultivar (Xiushui134) was reserved as an independent test accession. Whole-genome DNA sequences were segmented into 32 kb genomic regions using a sliding window with a step size of 16 kb, each such window was directly used as a context input for the model (i.e., 50% overlap between adjacent windows). Strand-specific RNA-seq reads were aligned to the corresponding cultivar reference genome. Reads mapped to chloroplast genomes were removed prior to downstream analysis. Base-resolution expression BigWig coverage tracks were generated using deepTools bamCoverage with a bin size of 1 bp and counts-per-million (CPM) normalization. Training and test chromosomes were partitioned prior to model fitting to prevent data leakage.

#### Model architecture

DNA encoder: The pretrained OGR foundation model (rice 1.25 B, 32 kb context) was loaded and served as the sequence embedding backbone for genomic representation learning.

Downstream prediction head: A U-Net-style encoder-decoder module with configurable projection dimension (1024), four downsampling blocks, and a bottleneck layer (1536) was appended to predict single-base expression tracks. Skip connections preserved fine-scale positional information during upsampling.

#### Training procedure

Full-parameter fine-tuning was performed in a distributed data-parallel (DDP) environment with synchronized batch normalization and bfloat16 mixed-precision computation. FlashAttention-2 was enabled to accelerate self-attention operations and improve computational efficiency. Model optimization was performed using the Adafactor optimizer with a cosine learning-rate decay schedule (base lr=1×10⁻⁴, 10% warmup), weight decay of 0.01, and gradient clipping (max norm=1.0). The model was trained for 20 epochs with a per-device batch size of 1, with gradient accumulation employed to accommodate GPU memory constraints. The loss function was mean squared error (MSE) between predicted and observed expression tracks. A custom Trainer module was implemented to support chromosome-aware sampling, per-track loss aggregation, and step-wise metric logging. Training dynamics and model checkpoints were monitored via Weights & Biases, with checkpoints retained based on minimum validation loss.

#### Evaluation

Model performance was assessed at both base-pair and gene levels. Agreement between predicted and observed expression values was quantified using Pearson correlation coefficient (PCC) computed on raw and log1p-transformed expression values, as well as the Spearman rank correlation coefficient. Performance metrics were computed independently for each chromosome to evaluate genome-wide generalization and then aggregated across genes to obtain overall model performance.

#### In silico variant effect prediction

To assess the regulatory impact of cis-acting sequence variants, we implemented an *in silico* mutagenesis framework. A 32,768 bp genomic context window encompassing the target locus was extracted from the reference genome. Mutant sequences were generated by introducing single-nucleotide substitutions at specified coordinates while preserving the surrounding sequence context. Both reference and mutant sequences were tokenized and processed through the fine-tuned model in evaluation mode with gradient computation disabled (torch.no_grad). The model outputs base-resolution RNA-seq prediction tracks for each sequence. Gene-level expression was quantified by summing the predicted signal across the annotated genomic coordinates of the target gene. As a proof of concept example, we applied this framework to *TAC1* (LOC_Os09g35980), evaluating a documented A-to-G substitution at the 3′-splice site of intron 4. This mutation disrupts canonical splicing and modulates tiller angle architecture, thereby demonstrating the fine-tuned OGR model’s capacity to link sequence-level perturbations to transcriptional and phenotypic outputs.

### Gene expression prediction based on multi-modal data

#### Datasets

The paired ATAC-seq and RNA-seq data were downloaded from SRA study SRP425619. Only shoot apical meristem tissue samples were used in this study. The reference genomes assemblies for MH63, ZS97, and Nipponbare (NIP) corresponded to GCA_016134885.1, GCA_001623345.3, and GCF_034140825.1, respectively. Although sequencing reads were processed genome-wide, only chromosome 3 was used for model training and evaluation. The training and test datasets were separated before model training to ensure that no test data were used during optimization.

#### Data preprocessing

ATAC-seq datasets: Paired-end ATAC libraries were processed using a standard analysis pipeline including adapter trimming, genome alignment, BAM file sorting, and coverage normalization. Trimmomatic was used for adapter trimming to obtain clean reads. The clean reads were mapped to the corresponding cultivar reference genome using Bowtie2. The resulting BAM file was sorted by coordinate, and reads mapped to the chloroplast were excluded. The BAM file was converted to BigWig format using deepTools bamCoverage, with a minimum mapping quality threshold of 30, a bin size of 5 bp, and CPM normalization.

RNA-seq datasets: Strand-specific RNA-seq reads were aligned and converted to separate plus-and minus-strand coverage tracks, CPM-normalized, at base-pair resolution. HISAT2 was used for RNA-seq alignment against the cultivar genome. The resulting BAM files were sorted by coordinate, and reads mapped to the chloroplast were excluded. The BAM files were converted to BigWig format using deepTools bamCoverage, with minimum mapping quality set to 0, and bin size of 1 bp, and CPM normalization, for both the plus and minus strands.

#### Model architecture

DNA encoder: The pretrained OGR foundation model (rice 1.25 B, 32-kb context) was used as the DNA sequence encoder. During fine-tuning, only the final transformer layer was trainable, whereas all preceding layers remain frozen.

ATAC encoder: ATAC-seq accessibility along the genomic window were encoded using a dedicated transformer tower. At each genomic position, scalar ATAC-seq accessibility values were lifted to a 192-dimensional embedding space through a linear layer. The resulting embedding were processed through six pre-norm transformer layers, each consisting of four-headed self-attention with a head dimension of 48, rotary positional embeddings matched to the DNA backbone’s RoPE settings, and a feed-forward sublayer whose hidden width is four times the model width (768). A final linear map projects the sequence from width 192 to width 1024 so the ATAC branch matches the DNA embedding dimension; the tensor is returned in channel-first form with length aligned to the DNA token grid.

Fusion and decoder: DNA and ATAC-seq representations were first downsampled to one-quarter of their original sequence length and subsequently fused with bidirectional cross-attention. The downsampled DNA and ATAC repesentations and two directional cross-attention features were then passed through a four-way gated MLP fusion module. A U-Net-style decoder restores full-length predictions, with skip connections from both DNA and ATAC branches for fine-scale reconstruction. Gate entropy regularization was applied at 1% of the MSE scale for both the fusion gates and the skip-connection gates, encouraging less peaked gating without dominating the regression objective.

#### Training procedure

Data Transform: The chromosome sequences were tiled with 32 kb windows and 16 kb overlap (50% overlap between consecutive windows). RNA expression levels were transformed before training using the method proposed by AlphaGenome (Avsec et al., 2026): consisting of a 0.75 power transform followed by squashing of extreme values. The inverse of this map was applied when comparing predictions to observations in the original signal domain.

Training Object: The training objective was defined as the sum of plus- and minus-strand mean squared error (MSE) between the predicted two-channel RNA expression tracks and the corresponding transformed targets signals. Additional entropy regularization terms proportional to the normalized Shannon entropy of fusion and skip-connection gate weights were incorporated at 1% of the MSE scale to discourage excessively peaked gating distributions.

Optimization: Model training was performed for 80 epochs using the Adafactor optimizer. Each optimization step used five windows per accelerator after twenty micro-batch accumulations, and accumulation was rescaled with the number of parallel processes so that the effective global batch size stayed comparable when more than one GPU was used. Discriminative learning rates were set to 3 × 10⁻⁵ on the trainable last DNA block and to 1.5 × 10⁻⁴ on all other trainable modules (ATAC encoder, fusion, and decoder head). The learning rate followed linear decay with 5% linear warm-up at the start of training. We used weight decay 0.01, gradient clipping at global L2 norm 1.0, and logged the training loss every optimizer step. Model weights were written to disk every five epochs while retaining at most ten checkpoints for downstream evaluation.

#### Evaluation

Agreement between predicted and observed RNA, pooled over bases, used the Pearson correlation coefficient and the coefficient of determination (*R*^2^).

## DATA AND CODE AVAILABILITY

The implementation details, model weights, and inference code for OneGenome-Rice are available at the GitHub repository: https://github.com/zhejianglab/OneGenome-Rice and https://github.com/BGI-Plant/OneGenome-Rice. The 26 benchmark datasets curated for rice genomics have been released on the Hugging Face Hub and ModelScope under the "RiceBenchmark" repository. Training sequences for the 422 accessions are accessible under the respective BioProject accession codes (Supplementary Table S1).

## ACKNOWLEDGMENTS

The authors thank Zhejiang Lab for developing the 021 Science Foundation Model. The model training process was conducted on the 021 Large Science Model, Zero2X open platform, and Nanhu Computing Framework.

## AUTHER CONTRIBUTIONS

All authors (B. Qian, C. Liang, C. Qin, C. Liu, C. Zhang, C. Xu, D. Li, G. Xue, H. He, H. He, H. Zhang, D. Chen, J. Xu, J. Zhang, J. Sun, J. Jiang, K. Xia, L. Zhong, L. Chen, L. Fan, L. Liu, L. Shang, M. Qing, Q. Li, R. Huang, S. Zhu, S. Ma, S. Liu, S. Zhang, S. Fu, T. Wei, X. Xu, X. Jia, X. Xu, Y. Jing, Y. Xu, Y. Bian, Y. Zhao, Y. Xue, Y. Guo, Z. Xiao, Z. Li, Z. Li, Z. Yue, Z. Deng) contributed to the design, training, and programming of the OneGenome-Rice model. C. Liang, D. Li, J. Zhang, L. Fan, S. Zhu, S. Fu, X. Xu, Y. Xu, Y. Xue, Z. Li additionally led the writing and revision of the manuscript. All authors read and approved the final version.

## DISCLOSURE OF USE OF AI-ASSISTED TOOLS INCLUDING GENERATIVE AI

In the preparation of this manuscript, an AI-assisted tool (Doubao, ChatGPT, Gemini) was utilized to support the optimization of academic writing structure (e.g., organizing the logical flow of the Methodology section, Abstract and designing the flowchart), as well as refine the expression of technical content. All content generated or optimized with the assistance of this tool underwent a thorough process of review, verification, and revision by the authors to ensure accuracy, academic rigor, and consistency with the study’s original findings.

## Supplementary figures and legends

**Supplementary Figure 1.**
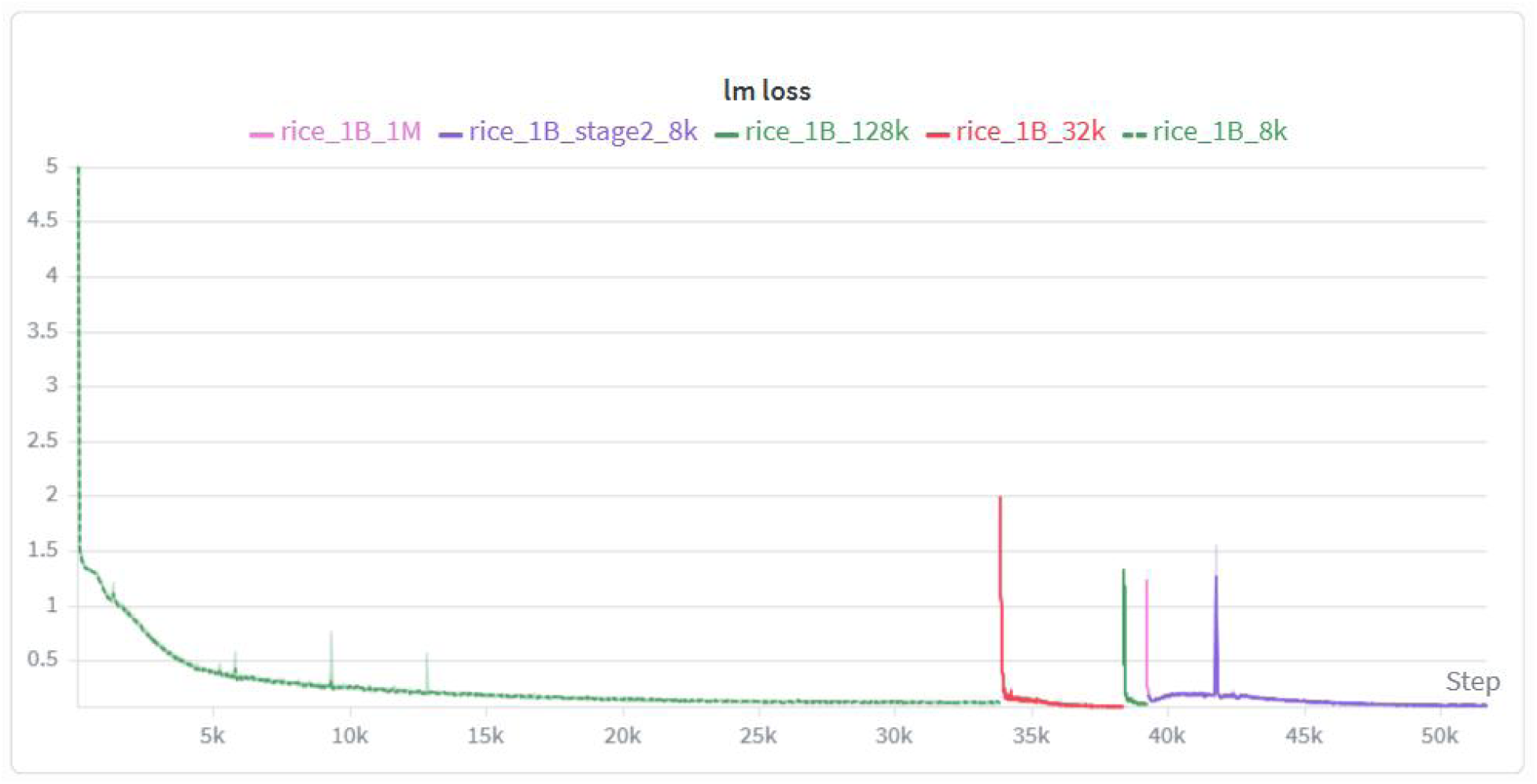
Modeling loss curve of OGR model based on Megatron framework in staged pre-training. This graph shows how the language modeling (LM) loss of the OGR model changes with the number of training steps during the pre-training phase. The x-axis represents the number of training steps (ranging from 0 to 52k), and the y-axis represents the LM loss value (ranging from 0.10 to 0.5). Each curve corresponds to a different OGR pre-training phase.

